# Electroencephalographic activity patterns distinguish interdigitated default and frontoparietal networks

**DOI:** 10.64898/2026.10.29.762898

**Authors:** Aaron Kucyi, David Braun, Lotus Shareef-Trudeau, Amir Adler, Dimitrios Pantazis

## Abstract

The brain is comprised of distributed functional networks that operate on diverse time scales. Evidence from precision fMRI shows that canonical networks, including default (DN), frontoparietal control (FPCN), and dorsal attention (dATN), each fractionate into multiple finely interdigitated networks. However, as fMRI is blind to fast processes underlying neural communication, there remains a need to neurophysiologically characterize individual-level network fractionation. To address this need, we conducted a simultaneous EEG-fMRI study, including multiple sessions per individual, resting-state and task conditions, and carbon wire loop sensors for optimized artifact correction. Using precision network mapping, we discovered dissociable electrophysiological correlates of blood-oxygen-level-dependent activity for two networks within the boundaries of the canonical DN. One (DNa) related more positively with alpha and high-beta power, while another (DNb) related more negatively with theta and low-beta power. Univariate and machine learning analyses further revealed electrophysiological dissociations between networks within the canonical FPCN (FPCNa versus FPCNb) but not dATN. The network-specific electrophysiological activity patterns were reliable across sessions, only modestly variable between resting-state and demanding task conditions, and observable for neighboring nodes of different networks across multiple cortical zones. Our findings provide insight into rhythmic neural processes that fine-grained networks may engage to support divergent cognitive operations.

## Introduction

The human brain is organized into a set of large-scale functional networks that operate over multiple time scales. Over the past two decades, functional connectivity analysis of fMRI data has remained the primary approach for mapping these networks during wakeful rest or cognitive task performance.^1,2^ A recently developed approach termed “precision fMRI,” or precision functional mapping, has placed emphasis on mapping the unique features of each individual’s functional networks.^3–6^ Careful mapping has led to the discoveries of person-specific features, and even entire networks, that were previously hidden from conventional group-level analyses.^7,8^ As a major example, canonical networks that are commonly studied in cognitive neuroscience, including default (DN), frontoparietal control (FPCN), and dorsal attention (dATN), each fractionate into at least two closely interdigitated networks that are distributed across multiple cortical zones.^9–13^ Such findings have important implications for how networks should be mapped and linked to function in research and clinical contexts.^6,14^ However, precision mapping methods critically rely on fMRI’s blood-oxygen-level-dependent (BOLD) signal, a slow hemodynamic measure that is not a direct readout of neural activity.^15^ There remains a critical need to corroborate individualized network estimates using neurophysiological measures that capture the fast time scales at which functional networks operate.

While precision fMRI research has predominantly focused on functional-anatomic features that differentiate brain networks, theoretical considerations and emerging empirical findings support the notion that closely interdigitated networks may be characterized by distinct temporal patterns of neural activity. Rhythmic activity has been proposed as a key mechanism that orchestrates long-range cortical communication throughout brain networks.^16–18^ Indeed, studies spanning multiple species and recording modalities suggest that different BOLD networks may arise from distinct patterns of spectral activity^6,19–26^ that could facilitate intra- and inter-network communication.^18,27^ In humans, extensive research using simultaneous EEG-fMRI has provided insight into electrophysiological correlates and predictors of BOLD fluctuations during rest and task states.^28–32^ Electrophysiological cofluctuations with network-level BOLD activity have most commonly been identified based on the amplitude of EEG band-limited power (BLP) at multiple frequencies.^33–43^ However, EEG-fMRI research has typically relied on the use of group-level templates, rather than individualized estimates, to define networks. As a potential consequence, inconsistencies have often been found across individuals, studies, and samples in EEG-BLP correlates of BOLD activity.^36,44^ For example, BOLD activity within regions of the canonical DN have been shown to relate positively, negatively, or neutrally with EEG alpha power.^33–37,39,45–47^

We suggest that the combination of simultaneous EEG-fMRI with the methodology of precision fMRI is well-suited for addressing key gaps of knowledge concerning the relationship between finely organized networks and temporal patterns of neural activity. Rich sampling of individuals, and person-specific mapping of BOLD networks, would permit rigorous tests of reliability and appropriately account for cross-subject variability in functional network anatomy. A finding of dissociable EEG spectral patterns for closely interdigitated networks could resolve prior inconsistencies regarding electrophysiological relationships with functional networks. Such a finding would also provide neurophysiological characterization of individual-level network fractionation and shed light on the role of fast neural processes in supporting divergent functions supported by different networks.

Here, we used simultaneous EEG-fMRI to investigate the electrophysiological bases of fine-grained networks identified with precision fMRI. Towards a “precision EEG-fMRI” approach, we applied multiple methodological advances that have not typically been used, or combined synergistically, in prior work. We leveraged a multi-session design, and we employed optimized correction of MRI-induced artifacts in EEG data using carbon wire loop sensors.^48^ In targeted analyses, we tested the hypothesis that EEG-BLP correlations are distinct for different personally-estimated subdivisions of the canonical DN (known as DNa and DNb^9^). In particular, given the selective role of DNa relative to DNb in mind-wandering, remembering the past, and imagining the future,^49–51^ we predicted that EEG features related to internally-guided modes of cognition (e.g. alpha power^52,53^) would more strongly track DNa than DNb BOLD activity. We subsequently sought to identify and detail the temporal relationships between EEG-BLP and fine-grained networks within the boundaries of the canonical DN, FPCN and dATN using both univariate and multivariate machine learning analyses. We target these specific canonical networks due their well-characterized fractionation within the individual.^9,10,49,54,55^ We further evaluate the degree to which network-specific EEG-BLP activity patterns are reliable across sessions, consistent between task and resting state conditions, and generalizable to the level of neighboring nodes of different networks across multiple cortical zones. Together, our findings offer electrophysiological evidence that enhances the interpretability of closely interdigitated brain networks defined within individuals.

## Results

### Precision functional mapping in a multisession simultaneous EEG-fMRI dataset

We acquired data from 23 healthy adult subjects who each underwent two simultaneous EEG-fMRI sessions, each on a different day.^56^ After censoring data points involving head motion, over 1 hour in total recordings of analyzable data were retained per subject (mean±SD = 73.2±7.7 minutes). During each session, subjects underwent two behavioral conditions, including a high-demand cognitive task requiring sustained attention and a minimal-demand resting state condition with intermittent sampling of thought contents.

In our EEG workflow (**Fig. 1a**), we preprocessed data from 31 scalp electrodes to remove non-neural noise due to MR gradients, and we regressed out signals from four carbon wire loop sensors to remove cardioballistic and helium pump artifacts (see Methods).^48^ We computed continuous BLP amplitude at frequencies ranging from 1-40 Hz. In selected analyses, the BLP estimates were averaged within canonical frequency bands (δ: 1-3 Hz, θ: 4-7 Hz, α: 8-12 Hz, β1: 13-20 Hz, β2: 21-30 Hz, γ: 31-40 Hz) and across channels. In our fMRI workflow (**Fig. 1b**), we performed precision mapping using multi-session hierarchical Bayesian modeling (MS-HBM)^55^ to estimate individual-specific functional anatomy of 17 cortical networks. Within each individual we identified six networks of interest, labeled according to Braga and Buckner,^9^ and extracted their preprocessed BOLD time series. The networks included parallel pairs of interdigitated networks (heretofore termed “parallel network pairs”) for the DN (DNa, DNb), FPCN (FPCNa, FPCNb), and dATN (dATNa, dATNb), as defined based on previously described criteria from studies that applied MS-HBM.^49,54,57^ Following EEG and fMRI processing, our main analyses focused on temporal relationships between EEG-BLP and BOLD activity averaged within each network (**Fig. 1c**). To maximize statistical power in our main analyses, we calculated relationships between EEG-BLP and BOLD activity using all available data per subject across sessions and behavioral conditions. As detailed further below, we also conducted analyses of reliability between sessions and variability between behavioral conditions (i.e., resting-state versus task).

**Figure 1.**
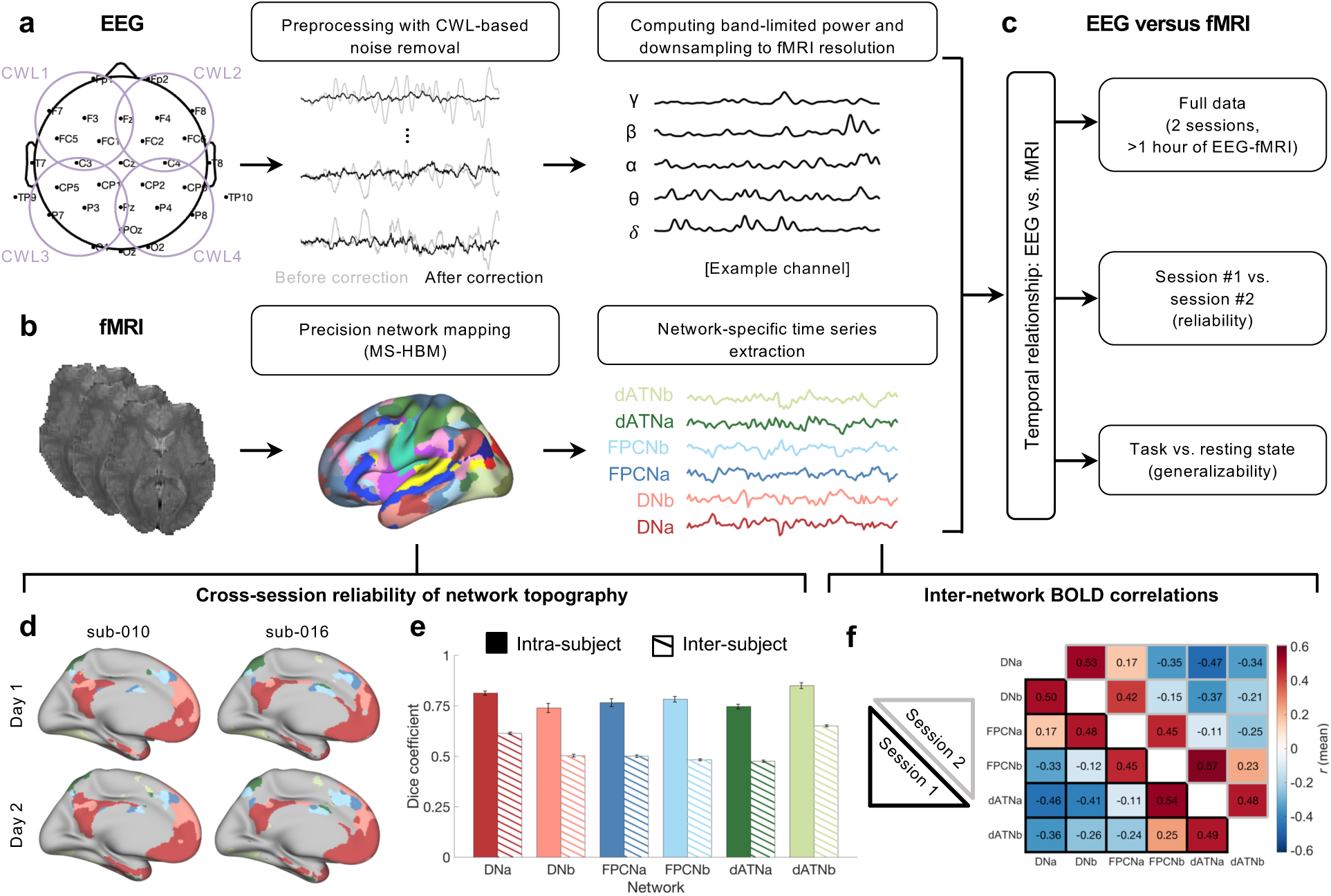
Analysis workflow for EEG-fMRI integration and precision network mapping. **a)** Scalp EEG data were acquired from 31 channels, while four carbon wire loop (CWL) sensors recorded magnetic field changes associated with movement. Following MRI gradient correction, a regression was applied to remove artifacts recorded by the CWL sensors. Band-limited power for frequencies ranging between 1-40 Hz was then computed, and averaging was performed within canonical bands (δ: 1-3 Hz, θ: 4-7 Hz, α: 8-12 Hz, β: 13-30 Hz, γ: 31-40 Hz; we also explored a split-up of β into β1 [13-20 Hz] and β2 [21-30 Hz]). **b)** Functional MRI data were preprocessed to minimize physiological and motion artifacts, and volumes acquired during high head motion were censored. Precision network mapping was performed using all fMRI data within each session, and time series from six networks of interest (DNa, DNb, FPCNa, FPCNb, dATNa, dATNb) were extracted. **c)** Temporal relationships were analyzed between EEG features (band-limited power) and network-specific BOLD time series. The relationships were analyzed in three different ways: (i) using all simultaneous EEG-fMRI data; (ii) using session #1 and #2 data separately; and (iii) using separate data for different behavioral conditions (resting state and Gradual-onset Continuous Performance Task; GradCPT). **d)** Example of networks obtained with precision mapping in two subjects, each at two different days (shown on medial surface). **e)** Group-level (n=23) mean dice coefficient values for six networks at the intra-subject level (session #1 versus #2) and inter-subject level (each session in a given subject versus all sessions in all other subjects). Error bars indicate standard error. **f)** Group-level (n=23) mean temporal correlation values for BOLD signals between all pairs of networks (bottom left triangle: session #1, top right triangle: session #2). Values are based on averages across all fMRI runs within a given session.

Prior to conducting main analyses, we verified that precision mapping yielded reliable network estimations and that the estimated networks showed expected patterns of BOLD functional connectivity within our data. Using MS-HBM based on fMRI data from session #1 and #2 respectively, we found that idiosyncratic features of networks were highly stable within individuals. Reproducible individual-specific functional-anatomic features were readily apparent in visualizations (**Fig. 1d**). For all six networks of interest, the Dice coefficient (a measure of spatial overlap between sessions) was substantially higher at the intra-subject (mean across networks: 0.78) relative to inter-subject (mean across networks: 0.54) level (**Fig. 1e**). Comparison of the individually-derived relative to population-based (i.e., Yeo17 atlas^58^) networks revealed substantially increased homogeneity of functional connectivity profiles for within-network vertices (**Fig. S1a**). The increased homogeneity was achieved due to considerable relabeling of population-based network vertices (**Fig. S1b**). Group-level inter-network correlations between BOLD signals in different individually-derived networks were highly stable across sessions (**Fig. 1f**). As expected, parallel network pairs showed positive BOLD correlations with one another. Across network pairs, the DNa and DNb showed negative BOLD correlations with all other networks except for FPCNa (in accordance with prior findings^59^). Despite these inter-network relationships, each single network carried unique variance which could potentially be temporally related to EEG-BLP fluctuations—a possibility that we explore next, first focusing on DNa versus DNb.

### Precision mapping refines relationships between EEG-BLP and BOLD activity in default network

In relating EEG-BLP to BOLD activity within fine-grained networks, our first aim was to replicate prior EEG-fMRI findings on electrophysiological correlates of the canonical DN. Positive correlations of DN activity with global (i.e., spatially-averaged across electrodes) EEG alpha power and negative correlations with delta/theta power have been previously reported^34,35,45^ (though see^33,36,37,39^). However, as prior works relied on group-average DN templates, it remains possible that these correlations have been based on blurred information across functionally heterogeneous regions within the boundaries of the canonical DN. To test this possibility, we compared EEG-BLP correlations with BOLD activity in the classical “Yeo7-DN” (based on the Yeo7 atlas^58^) (**Fig. 2a**) to correlations with BOLD activity in the individually-mapped DNa and DNb, which were both largely contained within the boundaries of the Yeo-7 DN (**Fig. 2b**).

**Figure 2.**
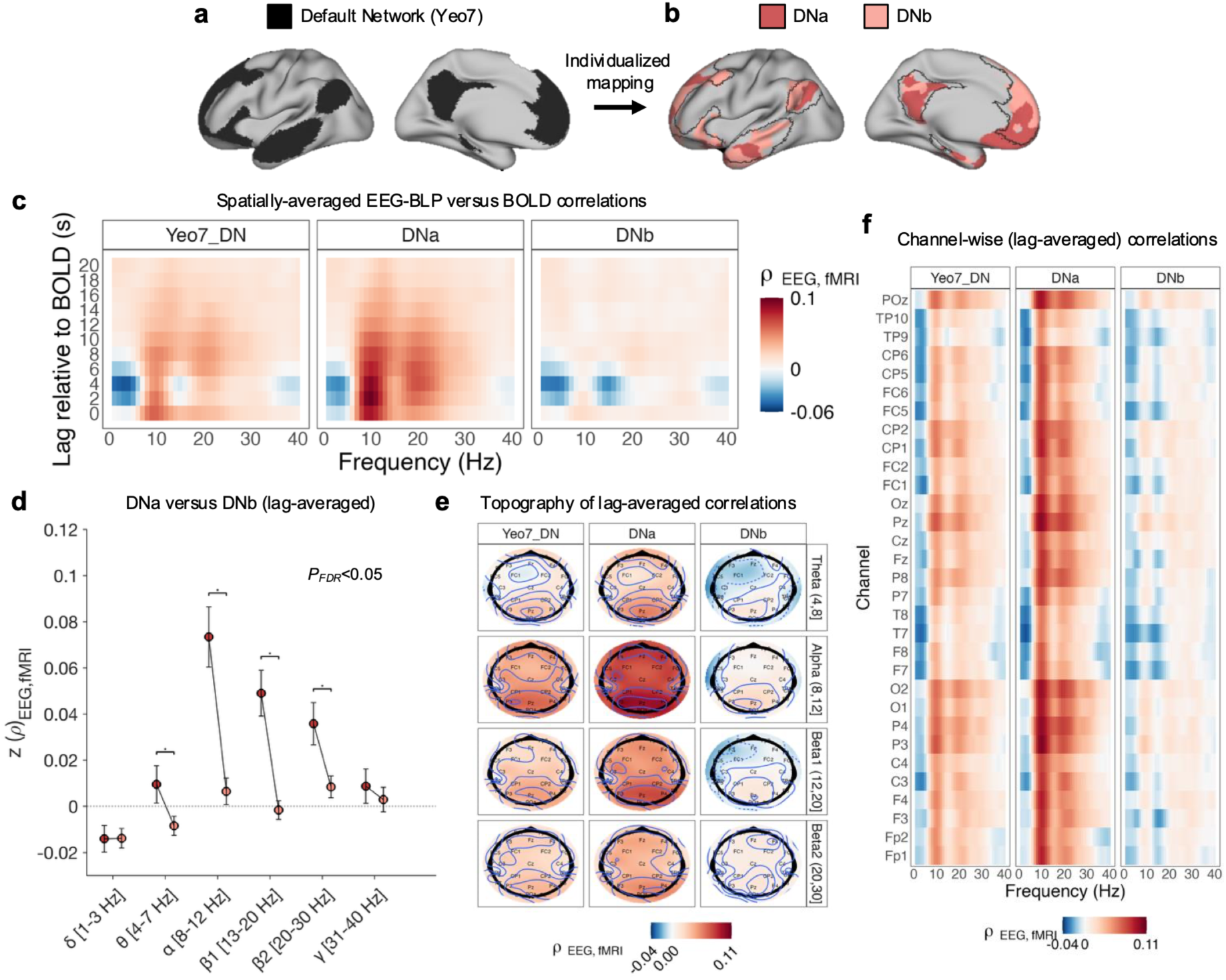
Dissociable EEG correlates of BOLD activity within DNa versus DNb. **a)** Functional anatomy of the canonical default network (DN) based on the standard Yeo7 atlas. **b)** Functional anatomy of DNa (red) versus DNb (pink) following precision functional mapping in an example subject. Black outlines indicate the boundaries of the canonical DN. **c)** Spearman rank correlation (ρ) between EEG band-limited power (EEG-BLP) and BOLD activity at multiple lag shifts of EEG relative to BOLD data (average based on all subjects and sessions). The EEG-BLP estimates were spatially averaged across all cortical electrodes. **d)** Comparisons between DNa (red dots) and DNb (pink dots) in terms of temporal correlation with EEG-BLP at different frequency bands (spatially-averaged across electrodes and time-averaged for lags between 0 to -10 seconds relative to BOLD sampling). Correlations are shown as Fisher z-transformed ρ values based on all subjects and sessions. Black lines/asterisks indicate significant differences between networks in terms of the strength of BOLD association with EEG-BLP (P_FDR_<0.05, Wilcoxon signed rank test). **e)** Topographic maps showing spatial variability in the correlation between BOLD activity and EEG-BLP at the frequency bands that showed significant differences among networks (theta, alpha, beta1 and beta2), based on averaging across all subjects and sessions (average for lags between 0 to -10 seconds). **f)** Channel-wise heatmaps showing spatial variability in the correlation between BOLD activity and EEG-BLP, based on averaging across all subjects and sessions (average for lags between 0 to -10 seconds).

For the Yeo7-DN, individually-mapped DNa, and individually-mapped DNb, respectively, we correlated global EEG-BLP versus BOLD activity at multiple lag shifts of EEG relative to BOLD ranging from 0 to -20 seconds (accounting for the sluggish and variable BOLD response to neural activity^15^). Correlations were conducted within each run, and then cross-run averaging was performed within subjects before group-level averages were computed. For the Yeo7-DN, we observed group-level positive correlations with EEG-BLP amplitude within the alpha range and negative correlations within the delta and theta ranges (**Fig 2c**, left), generally replicating prior findings.^45^ Importantly, the correlations were largest between 0 to -8 seconds of lag and were mostly absent at later lags. Given the well-known temporal characteristics of the hemodynamic response function,^60^ this observation is consistent with BOLD events being a direct consequence of EEG-related neural activity.

An inspection of group-level EEG-BLP correlation heatmaps for DNa (**Fig 2c**, middle) and DNb (**Fig 2c**, right) hinted that the correlations with Yeo7-DN were indeed likely confounded by spatial blurring of BOLD activity across DNa and DNb. Relative to the Yeo7-DN heatmap, the DNa heatmap showed stronger positive correlations with alpha and high-beta power at neurophysiologically relevant lags (i.e., peaking between -2 to -6 seconds). In contrast, for the DNb, positive correlations with alpha were largely absent, while negative correlations were present at lower frequency ranges and at the low-beta range. We statistically tested for differences between DNa and DNb in terms of magnitude of correlation with EEG-BLP within six frequency bands (temporally averaged between lags of 0 to -10 seconds) (**Fig. 2d**). A two-way repeated measures ANOVA revealed significant effects of frequency (*F*(5,110) = 15.3, *p* = 2.0x10^-11^) and network (*F*(1,22) = 24.6, *p* = 3.7x10^-5^) on correlation between EEG-BLP and BOLD activity. For theta, DNa showed significantly lower negative correlation than DNb (Wilcoxon sign-rank test, *p_FDR_ =* 0.0098). For alpha, beta1, and beta2, DNa showed significantly higher positive correlation than DNb (Wilcoxon sign-rank tests, respectively, *p_FDR_ =* 6.6x10^-5^*, p_FDR_ =* 1.0x10^-4^, and *p_FDR_ =* 1.9x10^-4^). No significant differences between DNa and DNb were found for delta or gamma frequencies. For EEG-BLP at the alpha frequency, which showed the largest between-network difference, 22 out of 23 individuals (95.7%) showed an effect that was congruent with the group-level finding of greater correlation with DNa than DNb. A null model analysis revealed that the differences between DNa and DNb exceeded those found when two surrogate networks with arbitrary boundaries, but preserved spatial interdigitation, were permuted with the territory comprising the true DNa and DNb (**Fig. S2**).

Our comparison of personally-mapped DNa and DNb to the Yeo7-DN template was motivated by the fact that prior EEG-fMRI studies largely treated the DN as a unitary network (i.e., as in the Yeo7 atlas). However, a 17-network, rather than 7-network, population-derived atlas (“Yeo17”) was used for spatial priors during our precision mapping procedures. We thus conducted a follow-up analysis in which we computed EEG-BLP correlations with BOLD activity in more granular, but population-derived, networks that were priors for DNa (“Yeo17-DNa”) and DNb (“Yeo17-DNb”). The EEG-BLP correlation heatmaps for Yeo17-DNa and Yeo17-DNb were largely similar, respectively, to the heatmaps for personally-mapped DNa and DNb (**Fig. S3a**). However, correlations with alpha and beta frequencies were stronger for DNa than Yeo17-DNa, suggesting refined EEG-BLP correlations when performing precision mapping (**Fig. S3b**). The amplified correlations with DNa were apparent within individual subjects who exhibited DNa and DNb spatial organization that deviated from population average patterns (**Fig. S3c**).

While the effects presented so far were based on global EEG-BLP, we further explored whether correlations were spatially heterogenous across different EEG electrodes. **Fig. 2e** displays topographic scalp maps of EEG-BLP correlations with Yeo7-DN, DNa, and DNb for frequencies at which DNa and DNb showed differences when electrodes were spatially averaged (i.e., theta, alpha, beta1 and beta2). At alpha, beta1, and beta2 frequencies, positive correlations with DNa were largely global but peaked at posterior occipital electrodes (**Fig. S4**). At theta and beta1 frequencies, negative correlations with DNb were widespread but peaked in left frontal regions. The topographic scalp maps were necessarily coarse due to the use of a 31-channel montage. Channel-wise heatmaps provided some further clarification of the largely spatially global nature of EEG-BLP correlations (**Fig. 2f**).

Taken together, these analyses revealed that the use of precision network mapping refines correlations between EEG-BLP and BOLD activity in DN that are otherwise obscured due to blurring together of DNa- and DNb-specific signals. We next extended our analyses to other parallel network pairs using both univariate and multivariate EEG-fMRI predictive modeling.

### Network-specific EEG-BLP patterns distinguish members of parallel network pairs

Precision network mapping has revealed parallel network pairs within the boundaries of the canonical DN, FPCN and dATN.^9,10,55^ Simultaneous EEG-fMRI studies have reported somewhat inconsistent EEG-BLP correlates of all the these networks across studies and datasets.^33–36,45,61^ Our analyses of DNa and DNb reported above highlight how such inconsistencies may stem from treatment of a network as unitary rather than fractionated. We thus next sought to clarify how EEG-BLP relationships with BOLD networks could depend more broadly on individual-specific network subdivisions. We conducted systematic analyses of EEG-BLP relationships with BOLD activity for six networks within three individually-mapped parallel network pairs (DNa/b, FPCNa/b, dATNa/b) (**Fig. 3a**) using a combination of mass-univariate and multivariate machine learning analyses.

**Figure 3.**
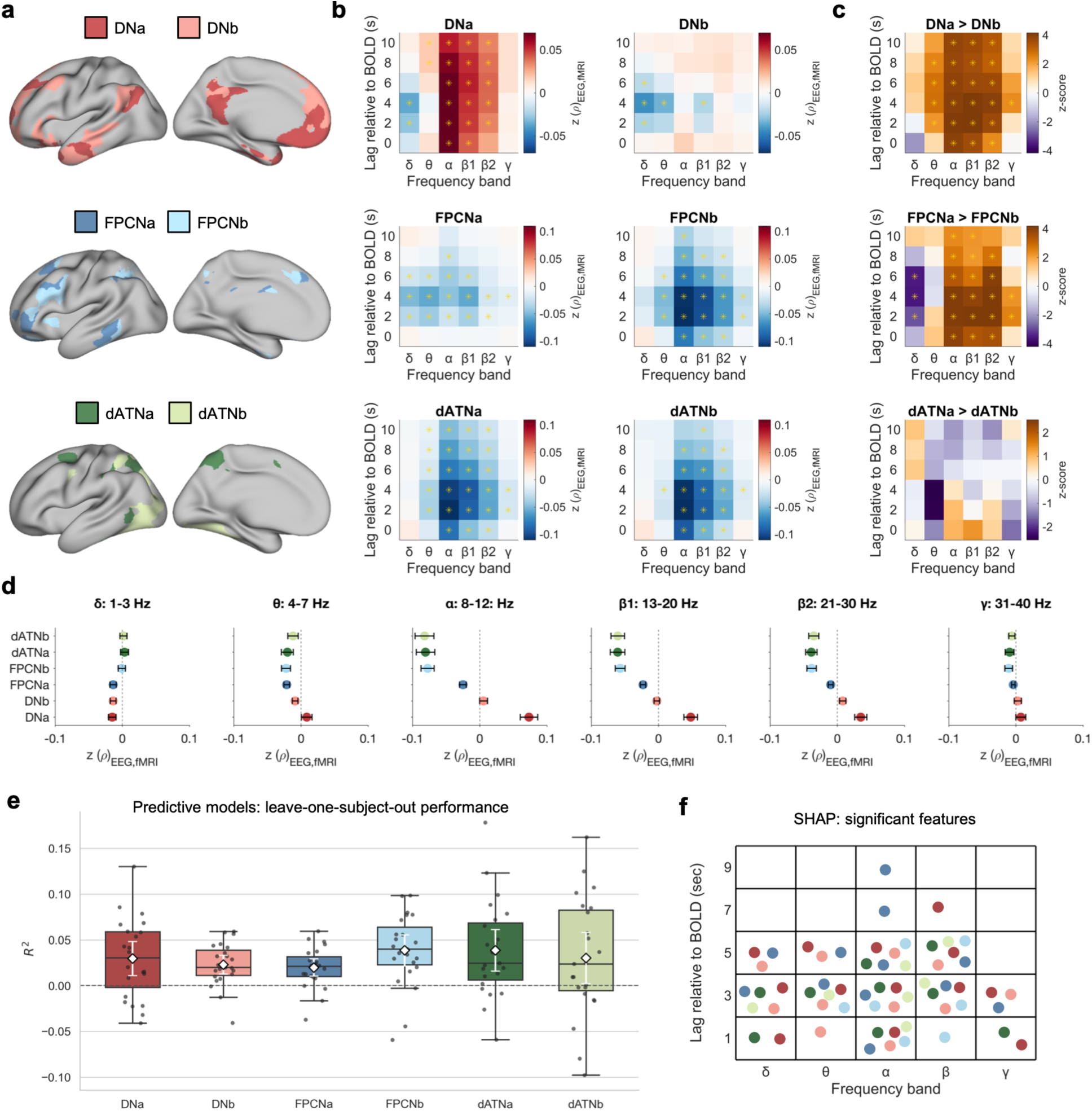
Network-specific EEG correlates of BOLD activity. **a)** Functional anatomy of parallel network pairs, including DNa (red) and DNb (pink), FPCNa (dark blue) and FPCNb (light blue), and dATNa (dark green) and dATNb (light green) shown on the cortical surface based on data from an example subject. **b)** Correlation heatmaps showing EEG-BLP versus BOLD activity for each network, at lags ranging from 0 to -10 seconds, and for multiple frequency bands (based on all subjects and sessions). Associations are displayed as Fisher z-transformed Spearman rank correlation (ρ). Asterisks indicate significant correlations at the group level (P_FDR_<0.05, sign test for zero median). **c)** Comparisons (z-scores) between members of each parallel network pair for all frequencies and lags. Asterisks indicate significant differences at the group level (P_FDR_<0.05, Wilcoxon signed rank test). **d)** Mean correlation, averaged across temporal lags (0 to -10 seconds), between EEG-BLP and BOLD activity (shown as Fisher z-transformed ρ) as broken down into six frequency bands. Error bars indicate standard error of the mean. **e)** Results of predictive modeling using XGBoost illustrating prediction of BOLD activity within each network from EEG features using a leave-one-subject-out (LOSO) cross-validation scheme. Individual points show the coefficient of determination (R^2^) for each held-out subject; the dashed horizontal line at R^2^=0 indicates chance-level performance. Boxplots summarize the distribution of subject-level R^2^ values (center line: median; box: interquartile range, IQR = 25th–75th percentiles; whiskers: 1.5×IQR). White diamonds indicate the mean R^2^ across subjects, with white error bars denoting the 95% confidence interval of the mean. **f)** Summary of features (organized by frequency band and lag relative BOLD) that showed significant contributions to predictive model performance, as determined based on absolute SHAP feature importance values (p_FDR_<0.05, 500 permutations).

For each network, we computed mass-univariate correlations of BOLD activity with a set of 32 global EEG-BLP features (derived from six frequency bands within the 1-40 Hz range and six lags of EEG relative to BOLD within the 0 to -10 second range). Largely reproducing the results reported above, the DNa showed significant positive correlations with alpha, beta1 and beta2 power at lags ranging between 0-10 seconds as well as significant negative correlations with delta power at lags of 2-4 seconds, while the DNb showed significant negative correlations with delta, theta, and beta1 power at lags ranging between 2-6 seconds (sign tests for zero median, *P_FDR_*<0.05; **Fig 3b**, top). The FPCNa showed significant negative correlations with delta, theta, alpha, beta1, beta2, and gamma power largely at delays between 2-6 seconds, while the FPCNb showed significant negative correlations with theta, alpha, beta1, beta2 and gamma power most consistently at delays between 2-6 seconds (sign tests for zero median, *P_FDR_*<0.05; **Fig 3b**, middle). The dATNa showed significant negative correlations with theta, alpha, beta1, beta2 and gamma power largely at delays between 2-10 seconds, while the dATNb showed significant negative correlations with the same frequency bands as dATNa at largely similar delays (sign tests for zero median, *P_FDR_*<0.05; **Fig 3b**, bottom). Similar to DNa and DNb effects (Fig. 2c), the EEG-BLP correlation patterns for FPCNa, FPCNb, dATNa, dATNb were also apparent when not restricting analyses to predefined frequency bands, and all networks showed relationships that temporally peaked at neurophysiologically relevant delays (i.e., ∼2-8 seconds) between EEG and BOLD sampling (**Fig. S5**).

To determine whether each member of a parallel network pair had a unique profile of correlation with EEG-BLP, we performed statistical contrasts between members of each pair. For the DN pairs, DNa relative to DNb showed significantly greater positive coupling with alpha and beta2 and significantly greater negative coupling with delta, while the DNb relative to DNa showed significantly greater negative coupling with theta, beta1, and gamma (Wilcoxon signed rank tests, *P_FDR_*<0.05; **Fig 3c**, top). For the FPCN pairs, FPCNa showed significantly greater negative coupling than FPCNb with delta, while FPCNb showed significantly greater negative coupling than FPCNa with alpha, beta1, beta2 and gamma (Wilcoxon signed rank tests, *P_FDR_*<0.05; **Fig 3c**, middle). However, no significant differences were found between dATNa and dATNb (**Fig 3c**, bottom). The results of these parallel network pair comparisons, and the network-specific correlation heatmap patterns for EEG-BLP versus BOLD, remained similar when preprocessing fMRI data using aCompCor^62^ instead of global signal regression (**Fig. S6**). To summarize how different EEG frequencies were related to each network in our mass-univariate analyses, we plotted mean correlations for each frequency band (**Fig 3d**). This highlighted several general patterns, such as—within the alpha to high beta range—a consistent progression across networks (DNa ◊ DNb ◊ FPCNa ◊ FPCNb and dATNs) from more positive to more negative correlations. Building on our mass-univariate analyses, we further conducted a predictive modeling analysis that could handle a high-dimensional EEG-BLP feature set (i.e., including data retained at the channel level, without spatial averaging) to predict BOLD activity in each network. We used the XGBoost algorithm^63^ to further account for potential nonlinear relationships and multivariate interactions across EEG-BLP features. Using a leave-one-subject-out cross-validation framework, we found that BOLD activity within all six networks was successfully predicted from EEG-BLP features at an above-chance significance level (DNa: mean R^2^ = 0.029, *p* = 2.7x10^-3^; DNb: mean R^2^ = 0.023, *p* = 9.1x10^-5^; FPCNa: mean R^2^ = 0.020, *p* = 2.0x10^-4^; FPCNb: mean R^2^ = 0.038, *p* = 2.4x10^-4^; dATNa: mean R^2^ = 0.039, *p* = 5.6x10^-4^; dATNb: mean R^2^ = 0.030, *p* = 1.9x10^-2^; **Fig. 3e**). We interpreted predictive models by computing Shapley additive explanation (SHAP) feature importance values with permutation tests to identify EEG-BLP features that contributed significantly to model performance for each network. Overall, frequency bands and lags that contributed significantly to performance (*p_FDR_*<0.05, 500 permutations) were largely consistent with mass-univariate results, but the predictive models also leveraged multivariate information from several frequency bands that did not show univariate effects (e.g. gamma contributed to DNa and DNb predictions but did not show univariate effects) (**Fig. 3f**). A description of predictive features, broken down by frequency band and temporal lag, is provided in **Fig. S7**. Predictive performance was driven by network-specific spatial distributions of EEG-BLP features (**Fig. S8**). Prediction of BOLD activity for each network, except for dATNb, was also successful when EEG spatial features were collapsed across channels; however, the effect sizes of prediction performance were decreased relative to when modeling with channel-wise features included (**Fig. S9**). This suggests that global EEG-BLP preserves predictive information (in line with univariate models) but that spatial EEG information carries additional variance relevant to network-level BOLD activity.

Overall, both mass-univariate and predictive modeling analyses confirmed that specific temporal patterns of EEG-BLP activity could be found for each network and that the members of DN and FPCN (but not dATN) parallel network pairs had distinguishable electrophysiological profiles at the level of global EEG signals. In subsequent analyses, we sought to determine whether the electrophysiological network patterns were reliable within individuals across sessions, stable across different task conditions, and generalizable to the level of neighboring nodes of different networks across multiple cortical zones.

### Network-specific EEG activity patterns are reliable across sessions

Results presented so far were based on combined data across two sessions (each occurring on a different day) per subject. To investigate reliability of the EEG-BLP correlations with BOLD activity in each network, we conducted analyses of independent data from each session. At the group level, side-by-side visual comparisons of session-specific EEG-fMRI correlation heatmaps revealed strikingly similar global patterns for each network at session #1 versus session #2 (**Fig. 4a**), with single-session results largely recapitulating the heatmaps that we had generated from the average across sessions (e.g. as shown in **Fig. 2c** and **Fig. 3b**).

**Figure 4.**
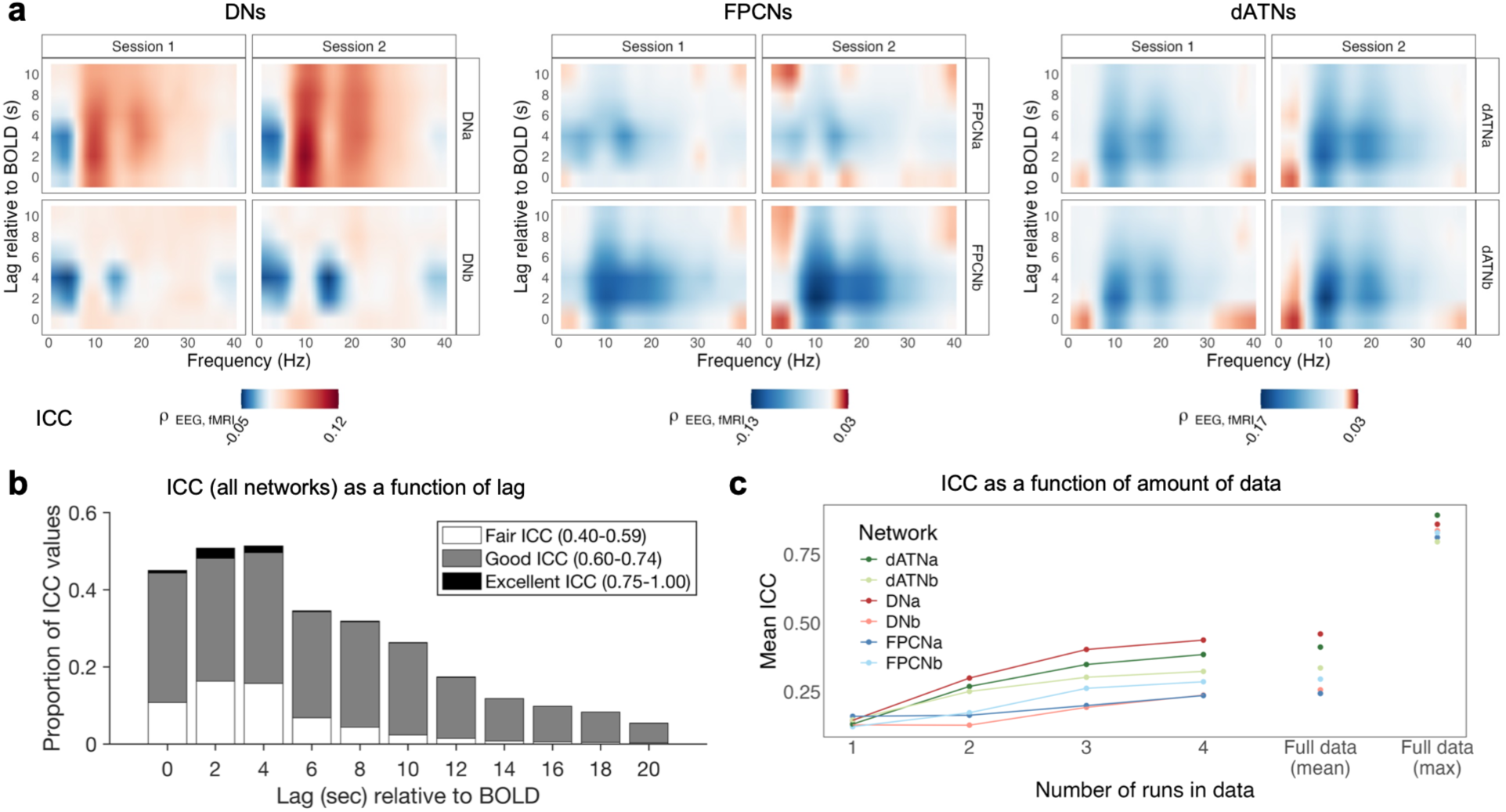
Cross-session reliability of correlations between EEG-BLP and network-specific BOLD activity. **a)** For each network (including parallel network pairs within DN, FPCN and dATN), correlation heatmaps are shown for session #1 and session #2 data separately (mean across 23 subjects). **b)** Proportion of intra-class correlation (ICC) values computed based on session #1 versus #2 data, that were within the “fair”, “good” and “excellent” ranges as a function of lag relative to BOLD (based on EEG-BLP between 1-40 Hz with ICC computed for each 1 Hz bin). **c)** Plot indicating how cross-session reliability varied with amount of data used to compute ICC within each session. For each network, the mean ICC (across all frequency bins and lags) increased when including up to 4 runs (∼8 minutes per run with runs trimmed to match duration across datasets) and further increased when using all available data per session (without trimming). For each network, the maximum ICC value (after screening all frequency-lag pairs and channels) was in the excellent range (>0.75).

To determine reliability within individuals, we computed the intraclass correlation (ICC) between sessions for multiple frequencies (1-Hz bins from 1-40 Hz) and lags of EEG-BLP relative to BOLD (2-second bins from 0 to -20 seconds) at each of the 31 EEG channels. The ICC values ranged from levels considered to be poor (<0.40) to excellent (>0.75) depending on the specific frequency-lag pair; this range was expected because only a small proportion among all EEG-BLP features were significantly correlated with BOLD activity within any given network. Importantly, features at neurophysiologically relevant lags of EEG relative to BOLD (especially -2 and -4 seconds) showed the highest proportion of ICC values within the fair, good and excellent ranges (**Fig. 4b**).

We further characterized how reliability varied with amount of data included within each session. As expected, the mean ICC (across all EEG channels and frequency-lag feature pairs) for each network increased when the number of included runs (∼8 minutes each; 4 runs per session) for ICC computation increased (**Fig. 4c**). Using all available data per session, we identified the maximum ICC value (ICC_max_) across all EEG-BLP features for each network. We found that ICC values considered to be in the excellent range could be identified for each network (DNa: ICC_max_= 0.86, DNb: ICC_max_=0.84, FPCNa: ICC_max_=0.81, FPCNb: ICC_max_=0.83, dATNa: ICC_max_ = 0.89, dATNb: ICC_max_ = 0.80). Taken together, our inter-session analyses revealed that excellent reliability of some correlations between EEG-BLP and BOLD activity could be identified for each network, especially at neurophysiologically relevant lags and when increasing the amount of data included. However, reliability was variable across lags, frequencies, and networks (**Fig. S10**).

### Network-specific EEG activity patterns vary modestly across task conditions

We have so far presented results that are based on EEG-fMRI data acquired during a combination of distinct tasks. Intrinsic activity within networks is persistent across both rest and task states, which explains why these networks can be readily identified from ongoing activity regardless of behavioral context.^64,65^ However, it is possible that the coupling between EEG-BLP and BOLD activity could depend on task-related or spontaneous cognitive processes.^37^ We acquired EEG-fMRI during conditions involving cognitive demand that was high (a prolonged task requiring sustained attention) or low (blocks of task-free, eyes-open rest). Frequent, behaviorally-relevant events in our high-demand task, known as the gradual-onset continuous performance task (GradCPT), are known to elicit BOLD activations and deactivations^66^ and neuronal population activity^67^ in DN, FPCN and dATN regions. We therefore sought to determine whether relationships between EEG-BLP and BOLD varied between GradCPT (i.e., a conceptual ‘treadmill test’ for cognition^68^) and Rest (i.e., spontaneous activity).

When limiting analyses to the GradCPT condition, correlation heatmaps for EEG-BLP versus BOLD within each network (**Fig. 5a**) showed many similarities to the previously described combined-condition heatmaps (**Fig 3b**). During the GradCPT, significant correlations between EEG-BLP and BOLD were identified for each network (sign tests for zero median, *p*_FDR_<0.05), including positive correlations of alpha and beta1 with DNa, negative correlations of delta and theta with DNb, and negative correlations of multiple frequencies with FPCNa, FPCNb, dATNa and dATNb. When limiting analyses to the Rest condition, correlations were generally weaker in magnitude but the heatmap showed many similarities to the combined-condition and GradCPT heatmaps (**Fig. 5b**). During Rest, significant correlations between EEG-BLP and BOLD were identified for all networks except for dATNa and dATNb (sign tests for zero median, *p*_FDR_<0.05), including positive correlation of alpha with DNa, negative correlation of delta and beta1 with DNb, and negative correlations of multiple frequencies with FPCNa and FPCNb (subthreshold negative correlations were observed for dATNa and dATNb but did not reach significance).

**Figure 5.**
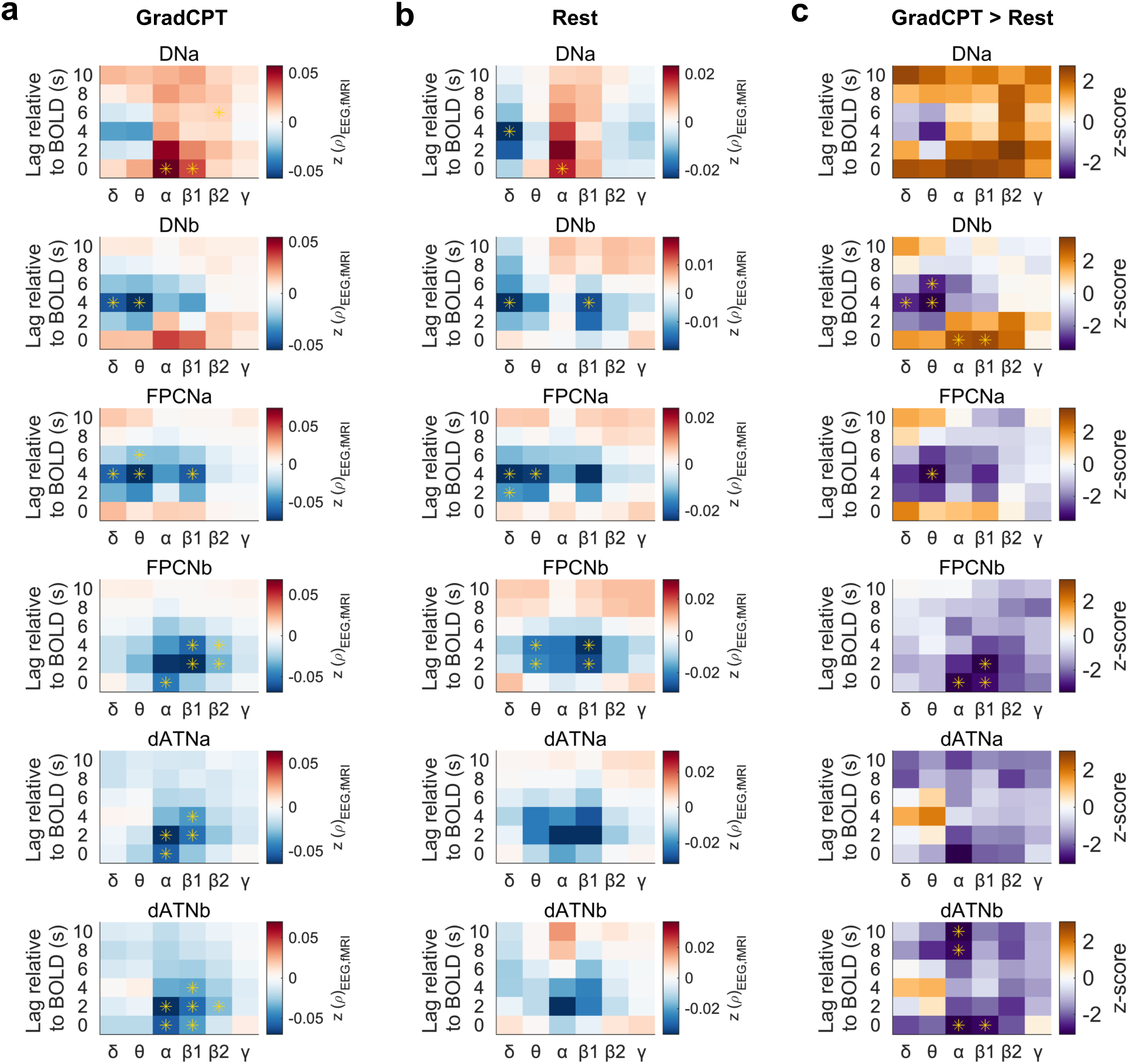
Comparison between task and resting state conditions. **a)** Heatmaps of correlation between BOLD activity in six networks and EEG-BLP (spatially-averaged across all EEG channels) at multiple frequencies and lags relative to BOLD. Correlations are shown as mean Fisher z-transformed Spearman ρ values across subjects. Asterisks indicate significant correlations at the group level (P_FDR_<0.05, sign test for zero median). **b)** Same as a) but for the Rest condition. **c)** Comparisons (z-scores) between GradCPT and Rest for each network. Asterisks indicate significant differences at the group level (P_FDR_<0.05, Wilcoxon signed rank test). All data shown are based on 20 subjects who had multisession GradCPT and Rest data.

A statistical comparison between GradCPT and Rest revealed significant differences in coupling between EEG-BLP and BOLD for four out of the six target networks (Wilcoxon signed rank tests, *P*_FDR_<0.05) (**Fig. 5c**). For DNb, there was greater positive correlation with alpha and beta1 as well as greater negative correlation with theta and delta for GradCPT relative to Rest. For FPCNa, there was greater negative correlation with theta for GradCPT relative to Rest. For FPCNb, there was greater negative correlation with alpha and beta1 for GradCPT relative to Rest. For dATNb, there was also greater negative correlation with alpha and beta1 for GradCPT relative to Rest.

In summary, absolute correlations (positive or negative) between EEG-BLP and BOLD were typically greater for the high-demand GradCPT compared to low-demand Rest condition. However, while the GradCPT may have strongly engaged the DN, FPCN, and dATN to drive greater EEG-BOLD coupling, the overall patterns of coupling at Rest were not drastically altered relative to GradCPT (despite the coupling being generally weaker). As such, the task-related variation in network-specific EEG correlates was modest and largely restricted to magnitude effects.

### Network-specific EEG activity patterns are observed at the level of neighboring nodes in different networks

We have so far reported results that were derived from mean BOLD signal within each network of interest. However, each network comprises a set of regions spanning multiple cortical zones. Within those zones, nodes of different networks (especially parallel network pairs) are often located in close spatial proximity to one another with shared boundaries. Do our identified EEG network-specific activity patterns generalize to side-by-side regions across distributed cortical areas? In other words, can distinctive EEG-BLP patterns be identified for BOLD activity extracted from different regions that are in close spatial proximity to one another?

To address this issue, we performed individualized fMRI areal parcellation^69^ of the cortex into 300 functional regions. We identified three sample cortical zones in inferior parietal lobule (IPL), medial prefrontal cortex (mPFC), and lateral temporal cortex (LTC) that contained neighboring regions that were members of DNa, DNb, and FCPNa (**Fig. 6a**; verified based on overlap with our individualized 17-network parcellations). For each zone (IPL, mPFC, LTC), we computed EEG-BLP versus BOLD correlation heatmaps (using the full dataset) for the neighboring regions. As can be seen in **Fig. 6b**, within each zone, the node-specific heatmaps showed patterns that largely resembled those found for the whole network that a given node belonged to (**Fig. 3b**). For example, positive correlations with alpha and beta1 were found for DNa nodes within IPL, mPFC and LTC. In contrast, positive correlations were absent for DNb nodes, which tended to instead show negative correlations with delta/theta and beta1 ranges. The FPCNa nodes that neighbored DNb and DNa tended to show negative correlations with a broader set of frequencies, including the alpha and beta1 range (i.e., an opposite pattern to DNa, even when a border was shared with a DNa node). As such, network-specific EEG activity patterns were largely generalizable to the level of neighboring nodes of different networks across multiple cortical zones.

**Figure 6.**
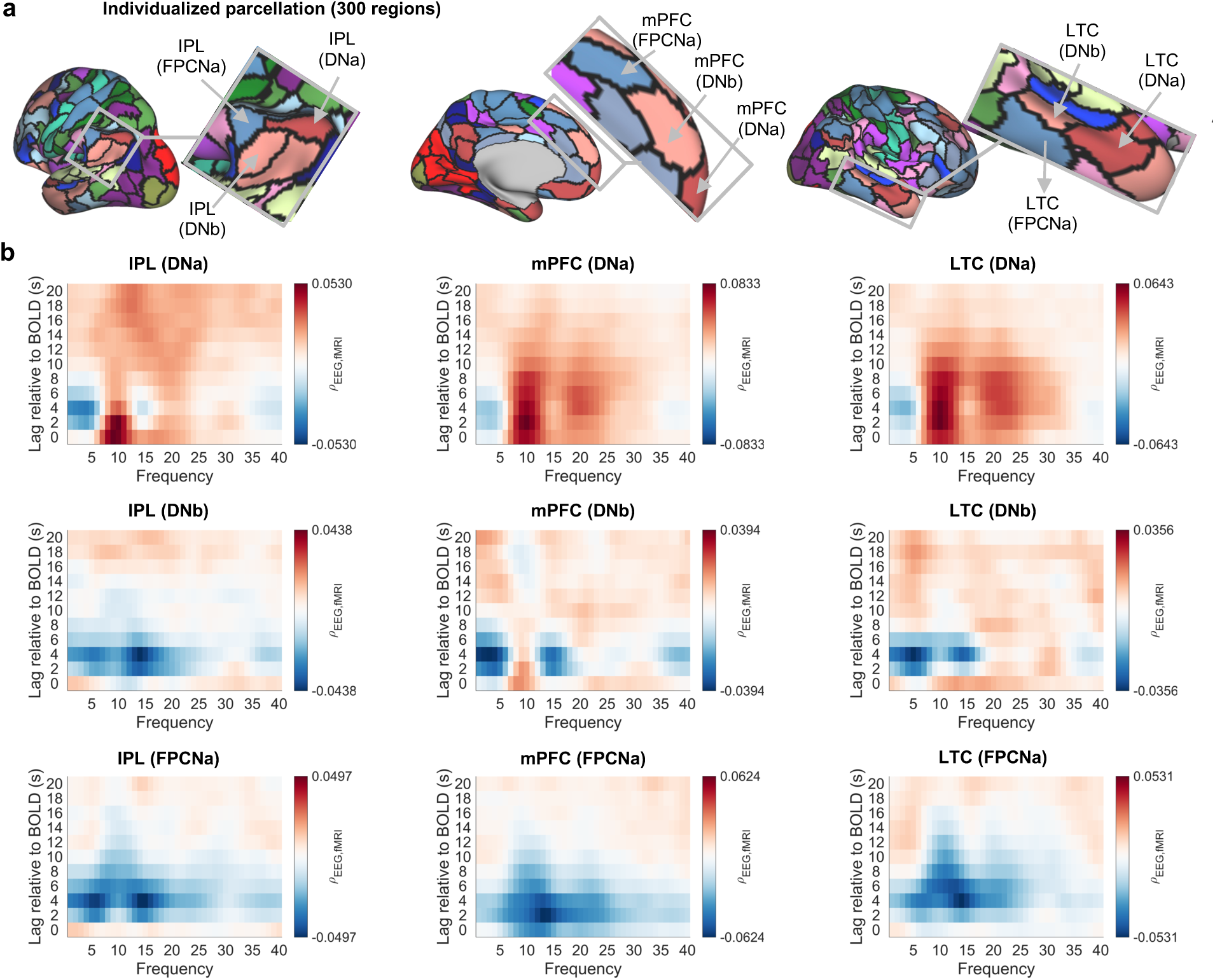
Relationships between EEG-BLP and BOLD activity for neighboring nodes of different networks across multiple cortical zones. **a)** Example of 300-region fMRI areal parcellation in a sample subject. Insets show close-up views of neighboring nodes that belong to different networks (DNa, DNb, FPCNa) within multiple cortical zones, including inferior parietal lobule (IPL), medial prefrontal cortex (mPFC), and lateral temporal cortex (LTC). **b)** Heatmaps of correlation between EEG-BLP and BOLD activity within each region highlighted in a). Correlations are shown as mean Spearman ρ values across subjects (n=23) for multiple frequencies and lags relative to BOLD (spatially-averaged across all EEG channels).

While our major analyses have focused on cortical regions, subcortical regions such as hippocampus have been associated with the canonical DN, and high-resolution precision fMRI studies suggest distinct affiliations of anterior versus posterior hippocampal subregions with DNa versus DNb.^10,12,70,71^ Although our fMRI sequences did not enable sufficient resolution to functionally map these subregions, we performed a preliminary analysis of EEG-BLP versus coarsely-defined anterior versus posterior hippocampus. This analysis revealed positive correlations between both hippocampal regions and EEG-BLP at alpha and high beta frequencies at neurophysiologically relevant lags (**Fig. S11**), resembling the DNa pattern (cf. Fig. 2c, 3b, and 4a). Thus, while our preliminary hippocampal analyses revealed limitations of extending our approaches to subcortical regions, they also provided electrophysiological support for the well-characterized relationship between the hippocampus and DN.

## Discussion

Using simultaneous EEG-fMRI combined with precision fMRI methodology, we identified reliable, frequency-specific, electrophysiological correlates of BOLD activity in individually-mapped functional brain networks. Distinct electrophysiological activity patterns were discernible for parallel network pairs within the boundaries of the canonical DN and FPCN. The DNa related more positively with alpha and high-beta power, while the DNb related more negatively with theta and low-beta power. Subdivisions of FPCN and dATN carried network-specific negative relationships with multiple electrophysiological frequencies ranging from delta to gamma ranges. The electrophysiological activity patterns for individual-specific networks were reliable across sessions, only modestly variable between resting state and demanding task conditions, and observable at the level of neighboring nodes of different networks across multiple cortical zones. Given the spatial specificity of the dissociations that we observed, our findings provide new knowledge that interdigitated network organization is not only predictive of cognitive processes (as shown in prior precision fMRI studies) but also of ongoing electrophysiological processes. Considered together with a growing body of neuroimaging research on the behavioral relevance individual-specific network organization, our results provide clues into how distinct, yet closely interdigitated, networks may engage fast neural processes to support cognitive function.

The DN is classically associated with deactivation during externally-oriented tasks and activation during internally-oriented mentation.^72^ Recent precision fMRI studies suggest preferential recruitment of DNa and DNb, respectively, during episodic memory recall and socially-oriented cognition.^50,73^ Despite these divergent functions, prior EEG-fMRI studies have largely treated the DN as a monolithic system or have not considered person-specific DN topography. Mixed findings have been reported regarding EEG correlates of the canonical DN, with some studies suggesting positive and others negative cofluctuations with alpha power.^33,34,36,37,39^ In a recent multi-dataset (albeit single-session per subject) analysis of resting state EEG-fMRI data, Xavier et al. consistently found negative delta-/theta-DN and positive alpha-DN correlations.^45^ Those EEG correlates of the canonical DN align more strongly with the activity patterns that we found for DNa rather than DNb. We speculate that prior inconsistencies in the literature may be explained by blurring of DNa and DNb signals as well as inclusion of regions that are not part of these networks within all individuals. Alternatively, future work involving more detailed, granular analyses of DNa and DNb subregions (and neighboring territories) may reveal heterogeneity in magnitude or sign of coupling with EEG-BLP.

Overall, our findings indicate that some of the advantage of individualized network mapping comes simply from considering the spatial scale of network definition, as EEG-BLP activity patterns distinguished DNa versus DNb even when these networks were defined in Yeo17^58^ population-based standard space. However, our findings further support a ‘sharpening’ of EEG-fMRI relationships through individualized network mapping; for example, DNa correlations with alpha and beta power were stronger when this network was personally-mapped rather defined based on a population-level atlas.

It has been theorized that alpha oscillations, which have long been known to increase during low task demands or eye closure (especially near occipital cortex),^74,75^ closely mirror the function of the DN.^37,76^ Our results provide support for a more specific coupling between alpha/beta and DNa, which may be in line with a gradient of function whereby regions in DNa, relative to DNb, are more insulated from externally-oriented inputs and outputs. Indeed, in densely sampled individuals, BOLD activation in DNa is more reliably associated than DNb with episodes of mind-wandering away from the external environment.^49^ Moreover, the DNa relative to DNb is more strongly connected functionally with the hippocampus (a finding supported by our analyses of hippocampal correlations with EEG-BLP),^10,71^ suggesting close cooperation with internally-oriented memory systems.

In contrast to the DN, the FPCN has been linked to functions such as executive control and top-down deliberative cognition.^77,78^ Regions within the canonical FPCN have extensive connections with both the DN and dATN and are thus well-positioned to mediate cross-network interaction and competition.^79^ A fractionation into FPCNa and FPCNb has been reported in group-level fMRI data,^58,59,80^ while precision functional mapping has further revealed individual variability in the topography of these two networks.^9,69^ The FPCNa and FPCNb have been shown to exhibit functional affiliation, respectively, with the DN and dATN.^59^ Our BOLD functional connectivity analyses produced results that align with those patterns: while FPCNa BOLD activity was positively correlated with both DNa and DNb and negatively with dATNa and dATNb, the inter-network coupling of the FPCNb showed inverse effects relative to FPCNa. The EEG patterns that we identified for FPCNa also more closely resembled the DNa and DNb patterns (e.g. included stronger negative correlation than FPCNb with delta band), while the FPCNb activity pattern was more similar to the dATN pattern (e.g. include stronger negative correlation than FPCNa with alpha/beta bands). As such, our findings offer neurophysiological support for previously documented preferential inter-network affiliations of the FPCNa and FPCNb. Early EEG-fMRI studies reported associations between alpha desynchronization and BOLD activity within canonical FPCN regions,^33,36^ but this effect was not consistently found in Xavier et al.’s recent cross-dataset analysis.^45^ Similar to effects that we have noted for the DN, the blurring together of FPCNa and FPCNb signals in group-level data provides a plausible explanation for such prior inconsistencies. Our data offer strong evidence for reliable EEG activity patterns that are dissociable for FPCNa versus FPCNb when these networks are mapped within individuals.

The dATN, associated with goal-directed cognition and visuospatial attention,^81^ exhibits an antagonistic (i.e., negatively correlated) relationship with the DN during task performance^66,67^ and, putatively, during resting states.^82,83^ This inter-network antagonism has been interpreted as a competition for access to shared computational resources.^84,85^ Several prior resting state EEG-fMRI studies have reported negative relationships between canonical dATN regions and alpha/beta power^33,34,45,61^ that are largely congruent with our findings. The negative coupling pattern that we found of alpha/beta with dATNa and dATNb is almost fully the inverse of the pattern that we found for DNa. In agreement with prior work that used group-level network templates,^86^ our findings suggest that the antagonistic relationship between dATN and DN regions may be mirrored in global electrophysiological states characterized by fluctuations in levels of neural synchronization at alpha/beta frequencies. At the level of global EEG signals, we did not find dissociable electrophysiological activity patterns for dATNa versus dATNb, and machine learning models revealed largely overlapping EEG features that explained BOLD activity in these two networks. Importantly, however, the lack of EEG-based dissociation between dATNa and dATNb does not necessarily imply that fMRI-based dATN fractionation is invalid. Empirical demonstrations of dissociable functions of dATNa versus dATNb have remained scarce, but anatomical layouts provide hints into potentially divergent roles in cognition (i.e., greater representation of intraparietal sulcus and frontal eye fields versus middle/lateral temporal visual areas, respectively, in dATNa versus dATNb). The split of dATN into two networks is highly replicated in individual-level fMRI data,^9,10,54^ and we confirmed both cross-session reliability and within-network homogeneity in our data. It is possible that a more sensitive approach, such as high-density EEG, is needed to characterize dATN fractionation in scalp-level electrophysiology. To our knowledge, this study represents the first effort to examine the within-individual reliability of temporal relationships between EEG-BLP and BOLD activity of functional networks. A cross-session comparison suggested a variable degree of reliability of associations between select EEG-BLP features and BOLD activity within each network investigated. While reliability improved with increased duration of EEG-fMRI recording, our design was not optimized for clarifying precisely how much data are needed to achieve maximum reliability. Future EEG-fMRI studies, ideally involving intensive sampling of individuals in longitudinal designs, are needed to more comprehensively assess reliability over time.

Our analyses cut across behavioral conditions involving cognitive demand that was either high (GradCPT) or low (resting state). However, most prior studies that have focused on continuous EEG relationships with BOLD activity have typically relied on resting state data alone (though see^40,42,87,88^). It has been suggested that coupling between alpha power and the DN is dependent on whether eyes are open or closed at rest,^37^ but broader EEG-fMRI comparisons of networks and frequency bands across various conditions have remained lacking. While alpha oscillations are a predominant electrophysiological pattern during wakeful rest, alpha power is also modulated by a wide variety of task conditions.^52,53,61,89,90^ Our results indicated an overall pattern whereby coupling between EEG-BLP and fMRI is relatively stronger during a high-demand task relative to rest (at least for DNs, FPCNs and dATNs). Prior work shows that during the GradCPT relative to rest, the magnitude of anticorrelation between regions of the DN and dATN (and other attention networks) is amplified.^67^ The sustained attention that is required for successful GradCPT performance is linked to frequent activations in dATNs and deactivations in DNs that may be associated with alpha desynchronization. At rest, spontaneous activations of these networks are expected to be decreased in frequency (but to still occur at irregular intervals), leading to weaker overall correlation strength with EEG-BLP fluctuations. Indeed, invasive electrophysiology studies have sometimes reported relatively weak correspondence between resting state BOLD signals and neural activity.^91–93^ As such, when the goal is to detect EEG predictors of fMRI networks, it may be desirable employ paradigms that explicitly evoke activations/deactivations, akin to ‘treadmill test’ for the networks of interest. However, further research is needed to determine the degree to which EEG-BLP coupling with BOLD is dependent on stages of cognitive processing or on sustained eyes-open versus-closed states that may differ in terms of alpha-DN coupling^37^ and behavioral role of alpha fluctuations.^89^

Importantly, dissociations that we observed within DNs and FPCNs were present when using channel-averaged (i.e., global) EEG metrics in our analyses. Fluctuations in physiological arousal that contribute to global changes in EEG-BLP and the fMRI signal have been well described.^94,95^ However, it is unlikely that our results can be explained by global arousal processes that unfold over a larger spatial scale than that captured with within-individual defined interdigitated networks. Indeed, a null model analysis (Figure S2) confirmed that the difference between DNa and DNb in associations with global EEG-BLP were spatially specific. It remains possible, however, that the associations were driven by differential engagement in arousal processes between these networks—a hypothesis that may be tested in future work that incorporates physiological measures.

Our findings have implications for the potential of using EEG alone to serve as a proxy of activity in individual-specific fMRI networks. As precision fMRI approaches typically involve high subject burden and cost, the alternative of EEG (including portable systems) could facilitate widened applicability in clinical contexts. The small correlation magnitudes that we observed were largely on par with effects previously reported.^34,45,86^ While our machine learning analyses revealed that within-network BOLD activity could be predicted from EEG-BLP features, prediction performance was typically low, suggesting that further refinements would likely be needed before EEG could be used to reliably index network-level BOLD activity. Prior work suggests that EEG can approximate BOLD activity in focal regions such as the amygdala^40^ but, in doing so, also targets wider brain networks.^96^ Our analyses also indicate that global EEG correlates of BOLD activity at single nodes of a given network often resemble the correlates of the whole network. It is possible, however, that high-density EEG combined with source localization techniques could be used to clarify contributions of EEG volume conduction^75^ and determine the spatial co-localization between EEG signals and fMRI networks.

The paradigm that we have described provides a blueprint for future work that may extend beyond our limited scope that centered on DNs, FPCNs, and dATNs. Important targets include additional cortical systems, such as sensory, motor-related and salience networks, as well as nodes within subcortical and cerebellar regions. Ultrahigh field fMRI with simultaneous EEG^97^ is needed to resolve relationships between hippocampal subregions and EEG-BLP. The thalamus represents an important region for future investigation due its role in generating and maintaining oscillatory neural activity.^75,90,98^ Recent evidence indicates that the thalamus can be functionally parcellated into network-specific subdivisions using precision fMRI.^64^ Improved understanding of broader electrophysiological relationships with BOLD activity could guide approaches for identifying causal relationships between rhythmic neural processes and individual-specific networks, such as use of neurostimulation or computational modeling.

In conclusion, our electrophysiological characterizations of precision fMRI provide key insights into neural processes that support subdivisions between fine-grained networks. Our multisession design and individualized approach to network definition allowed us to discover reliable and specific relationships between EEG-BLP and BOLD activity. Future work that may leverage this precision EEG-fMRI approach to shed further light on the electrophysiological bases of functional networks in individuals who are healthy or affected by illness.

## Materials and Methods

### Participants

Participants were 23 healthy adults (age range: 18-35; 14 female, 9 male) from a simultaneous EEG-fMRI dataset that we have made publicly available.^56^ While the original dataset included 24 subjects, one subject was excluded from the present analyses because they completed only one simultaneous EEG-fMRI session. Each included participant completed three total visits, including two that included simultaneous EEG-fMRI scanning. All participants had normal or corrected-to-normal vision, were right-handed, fluent in English, and reported no current or history of psychiatric, neurological, or chronic medical conditions. Participants provided informed consent for procedures approved by the Drexel University Institutional Review Board, and all methods reported here are in accordance with the approved protocol. Details on the full procedure of this study are reported elsewhere.^56^

### Neuroimaging data acquisition

Neuroimaging was performed on two different days for each participant with a 3-Tesla Siemens MAGNETOM Prisma scanner equipped with a 64-channel head coil. The T2* BOLD fMRI runs were performed with a CMRR multiband gradient-echo EPI single shot sequence aligned with the AC-PC plane (repetition time/TR: 2.0 sec, echo time: 25 ms, flip angle: 70°, slices: 81, field of view: 192mm x 192mm, voxel size = 2 x 2 x 2 mm). Within each session, subjects completed two tasks, including one run of the Gradual-Onset Continuous Performance Task (GradCPT; 8 minutes each) and three runs of resting state with experience sampling (average of 10.5 minutes each). At the beginning of the first neuroimaging session, a high resolution MPRAGE T1-weighted anatomical image was also acquired (TR = 2400 ms; TI = 1000 ms; TE = 2.22 ms; slices = 208; FoV = 256 x 240mm, voxel size = 0.8 mm^3^ isotropic; flip angle = 8°; partial Fourier off; pixel bandwidth = 220Hz/Px). At each session, a B0 field map was acquired (TR = 789 ms; TE 1 = 4.92 ms; TE 2 = 7.38 ms; slices = 81; FoV = 192mm x 192mm; voxel size = 2.0 mm^3^ isotropic; flip angle = 45°; partial Fourier off; pixel bandwidth = 668 Hz/Px).

### EEG data acquisition

Simultaneous EEG data were acquired during fMRI scanning using a 32-channel Brain Products MR-compatible system (Brain Products Gmbh, Gilching, Germany). Recorded channels included 4 carbon wire loop (CWL) sensors, 31 cortical channels and one electrocardiography channel placed on the back. Electrode impedance was kept below 20 kOhms during data acquisition, and the data were recorded at a sampling rate of 5kHz. Electrodes were arranged according to the international 10-20 system and included a reference and ground electrode. The CWL channels recorded head movements, allowing for optimized scanner artifact correction.^48,99^ A Brain Products SyncBox was used to synchronize the EEG phase with the MR clock. A Brain Products Triggerbox was used to send and receive MRI and stimulus triggers to EEG recording software.

### Tasks performed

The tasks performed during simultaneous EEG-fMRI were the GradCPT^66^ and a resting state with multidimensional experience sampling (rs-MDES) paradigm that we have previously reported.^100^ Briefly, during the GradCPT, participants observed a continuous stream of city or mountain scenes with gradual transition from one image to the next (800 ms total duration of each instance of each scene). Participants were instructed to press a button each time a city scene appeared (∼90% of trials) and withhold a press each time a mountain scene appeared (∼10% of trials). Behavioral data indicated that participants engaged in this high-demand task appropriately, as reaction time and accuracy performance data were within reasonable ranges.^56^

During the rs-MDES task, participants viewed a central fixation cross in 45-60 second intervals that were interleaved with experience sampling periods. For visual fixation periods, participants were instructed to keep their eyes open and focused on the central cross image while allowing their minds wander freely. During each experience sampling period, a set of 13 thought probes was presented, and subjects submitted ratings to indicate subjective qualities that they experienced prior the onset of the probe set. The experience sampling ratings were obtained for exploratory purposes and are not used further in the analyses presented. Stimuli for all task were presented using Psychophysics Toolbox.^101^ The detailed stimulation parameters and timing information are reported elsewhere.^56^

### fMRI preprocessing

The fMRI data were preprocessed with methods that were consistent with pipelines used previously to validate the MS-HBM approaches for precision network mapping.^55,69^ We used publicly available code from Computational Brain Imaging Group (https://github.com/ThomasYeoLab/CBIG/tree/master/stable_projects/preprocessing/CBIG_fMRI_Preproc2016). Structural data (T1 scan from first session) were reconstructed into surface mesh representations using the recon-all command via Freesurfer v7.4.1 software.^102^ Motion correction was performed using rigid body translation and rotation in FSL, and volumes containing a framewise displacement of >0.3 mm were marked for motion censoring. Spatial distortion correction was applied using FMRIB’s Utility for Geometrically Unwarping EPIs (FUGUE) tool with magnitude and phase-difference fieldmap images as inputs. Structural and functional images were aligned using boundary-based registration (using T1 structural images acquired in the first session).^103^ We then performed linear detrending and regressed out 9 regressors plus their temporal derivatives, including global signal, averaged WM signal, averaged CSF signal, and six motion correction parameters. A temporal bandpass filter (0.01 – 0.1 Hz) was then applied before fMRI data were projected to the Freesurfer fsaverage6 surface mesh. Spatial smoothing was performed on the surface with a 6mm full-width half-maximum kernel. Notably, the MS-HBM network mapping procedures were performed based on data that were temporally band-pass filtered (which was performed prior to surface projection). However, to preserve higher frequencies in the BOLD signal for integrated EEG-fMRI analyses, only a 0.01 Hz high-pass filter was applied when extracting BOLD time series.

While our main analyses involved cortical surface-based preprocessing, we also performed preprocessing in volumetric space to enable preliminary analyses of subcortical regions. The volumetric pipeline included the same preprocessing steps as the surface-based pipeline up to temporal filtering (0.01 Hz high-pass filter). After filtering, the native fMRI data were registered to MNI152 space via a series of steps using Freesurfer’s *mri_vol2vol* command where T1 anatomical scans were first projected to 1mm MNI152 space, then fMRI data were projected to 1mm MNI152 space, then the projected data were downsampled to 2mm MNI152 space. Spatial smoothing was then applied with a 6mm full-width half-maximum kernel.

Importantly, while our main fMRI preprocessing pipeline includes the step of global signal regression (GSR), it has been suggested that GSR introduces or inflates anticorrelations between the DN and other networks such as dATN.^104^ As such, we repeated several analyses based on fMRI data preprocessed with an alternative method, aCompCor,^62^ instead of GSR. In this alternative pipeline, the GSR step was replaced with aCompCor within the Computational Brain Imaging Group function “*CBIG_preproc_regression.csh*.” The top 5 principal components generated from a merged mask of white matter and cerebrospinal fluid regions were regressed out. The white matter and cerebrospinal fluid masks were eroded within T1 anatomical space prior to merging to avoid partial volumes that include gray matter. All other preprocessing steps carried out for the aCompCor pipeline were otherwise identical to those done for the GSR pipeline (a 0.01 Hz high-pass, not bandpass, filter was applied).

### Precision fMRI network mapping

We applied the MS-HBM method to parcellate each individual’s brain into 17 functional networks. We selected this method because MS-HBM has been extensively validated against several competing methods for precision functional mapping^55^ and has shown good correspondence with network parcellations obtained from other commonly used precision fMRI methods.^105^ For each subject, the MS-HBM procedures were applied separately to all fMRI data in sessions #1 versus #2 to enable tests of within-subject reliability. We submitted all fMRI data within a given session (i.e., all GradCPT and rs-MDES runs) to MS-HBM given that it is well established that task-related variance does not substantially alter the topography of individual-level parcellations.^64^

The detailed MS-HBM procedures are reported elsewhere.^69^ Briefly, in our implementation, we used pre-trained group priors that comprised 17 unique network labels for each vertex on the fsaverage6 surface. For each subject (and each session separately), functional connectivity profiles were computed (with exclusion of censored volumes), and each vertex was assigned to a network label such that the cerebral cortex was parcellated into 17 person-specific networks. Six networks of interest (DNa, DNb, FPCNa, FPCNb, dATNa, dATNb) were identified based on previously described criteria from studies that described these networks^9,10^ and applied MS-HBM to define them.^49,57,64^

Considering the inconsistency in network labeling schemes across prior fMRI studies,^106^ we manually labeled each network based on its functional anatomy to facilitate interpretability and compatibility with our prior work.^49,107^ As done previously,^49,107^ we did not strictly adhere to the Schaefer^108^ and Yeo^58^ labeling schemes that are provided within the MS-HBM pipeline. Our labels for the DNa, DNb, FPCNa, FPCNb, dATNa, and dATNb were consistent with Braga and Buckner^9^ who initially described fractionation of each of these networks into “a” and “b” networks within the individual. We acknowledge inconsistencies in the literature on the labeling of these networks; for example, the “A” and “B” labels for the “frontoparietal network” (here termed FPCN) have sometimes been flipped across different studies.^9,54,55,57,59,64^

Within DNs, the major zones containing interdigitated regions assigned to DNa and DNb were located within the posteromedial cortex, medial prefrontal cortex (with greater ventromedial representation for DNa^9,57^), lateral temporal lobe, medial temporal lobe and posterior parietal cortex (largely near angular gyrus). Within the FPCNs, the major zones containing interdigitated FPCNa and FPCNb regions were located within dorsolateral prefrontal cortex, ventrolateral prefrontal cortex (with greater rostrolateral representation for FCPNa^59^), inferior parietal cortex, inferior lateral temporal cortex, dorsomedial prefrontal cortex (near supplementary motor area), and anterior insular cortex. Within the dATNs, the major zones were located within dorsal frontal cortex (near frontal eye fields), superior parietal lobule, and lateral occipital cortex (with more posterior representation for dATNb^9,57^).

To determine inter-session reliability of the six networks, we computed the Dice coefficient as an index of intra-subject overlap of parcellations between sessions, as done previously.^69^ We computed a single Dice coefficient for each network and subject. We conducted a comparison to inter-subject overlap of parcellations, which was expected to be substantially lower than intra-subject overlap. To compute inter-subject values, for each subject and session, we computed the Dice coefficient based on a pairing with every other subject (each session independently). We then averaged the Dice coefficient values separately for intra-subject versus inter-subject levels. As a further comparison of fMRI data obtained independently at the two sessions, we computed functional connectivity (i.e., correlation) between all network pairs using preprocessed BOLD time courses, averaged across all runs within a given session.

To evaluate the quality of MS-HBM outputs, we performed comparisons with networks derived from the population-based Yeo 17-network atlas (Yeo17).^58^ Specifically, following previously reported methods, we computed “connectional homogeneity” within each network for MS-HBM-versus Yeo17-derived networks.^55,108^ We computed a homogeneity metric for each network by averaging the Pearson’s correlations between BOLD time series of all within-network vertices (averaged across all BOLD runs). We then averaged the correlations across all networks (separately for MS-HBM and Yeo-17 derived networks) while accounting for network size.^108^ Moreover, to illustrate how MS-HBM and Yeo17 networks diverge from one another, for each MS-HBM-derived network of interest, we computed the proportions of vertices that overlapped with each Yeo17 network. The proportions were averaged across sessions within subjects and then averaged across subjects.

In addition to our main analyses, which focused on mean BOLD activity within whole networks of interest, we further investigated single nodes of networks located within multiple cortical zones. To do so, we performed areal MS-HBM^69^ to define 300 individual-specific regions based on vertex-wise functional connectivity profiles. Similar to the 17-network MS-HBM procedure, we performed areal MS-HBM to all fMRI data in each session separately. The areal parcellations were performed using the gradient-infused MS-HBM approach (default parameters) and was guided by a group prior derived from fMRI data in 40 subjects in the Human Connectome Project.^69^ After performing areal MS-HBM, we selected three sample cortical zones for further analysis. These zones included the left inferior parietal lobule, medial prefrontal cortex, and lateral temporal cortex. We selected these zones based on inclusion of neighboring regions within DNa, DNb, and FCPNa (as verified based on visual inspections of 17-network parcellations). We extracted the preprocessed BOLD time series from each of these regions for submission to further analyses (i.e., testing associations with EEG features, as described below).

We also performed a preliminary analysis of BOLD activity within coarsely-defined hippocampal subregions. In volumetric MNI152 space, we defined the hippocampus based on the FSL Harvard-Oxford subcortical atlas, thresholded at 50% probability.^109^ We then split the hippocampus along the coronal plane at the y=-18 position to define anterior and posterior subdivisions. This position corresponds coarsely to the position of the uncal apex^110^ with a slight anterior shift to account for findings suggesting that the most anterior regions of medial temporal lobe may include more spatial representation of DNb than DNa.^70^

### EEG preprocessing

Preprocessing for all EEG data was run in MATLAB R2024b using EEGLABv2024.2.1.^111^ The EEG data were downsampled to 250Hz. We used specialized procedures for correction of scanner-induced artifacts in the EEG data. First, MR gradient induced artifacts were corrected using the BERGEN toolbox,^112^ which uses fMRI realignment parameters to subtract artifact templates from the EEG data. Next, cardioballistic and helium pump artifacts were corrected using CWL regression.^48^ Following scanner artifact correction, the data were bandpass filtered between 1 and 50 Hz. The data then were re-referenced using the average of all electrodes. We ran independent components analysis (ICA) to identify artifacts and rejected components using the Multiple Artifact Rejection Algorithm plugin.^113,114^ The data were then manually inspected for remaining artifacts and noisy channels to interpolate. Where interpolation was necessary, interpolation was applied to flagged channels in the pre-ICA EEG pipeline. We then re-ran ICA identification and rejection on the interpolated data, and any remaining artifacts identified manually were rejected.

### Univariate EEG-fMRI correlation analyses

To examine relationships between EEG features and BOLD activity, we first focused on EEG band-limited power (BLP) and fMRI default network (DN) activity. Within each run, BOLD time series were extracted from preprocessed fMRI data based separately on the mean of vertices within the “Yeo7-DN” derived from the standard-space Yeo7 atlas,^58^ the mean of vertices within the MS-HBM-derived DNa, and the mean of vertices within the MS-HBM-derived DNb. For further comparison of the MS-HBM-derived DNa and DNb to population-derived priors that were used during MS-HBM, we also extracted BOLD time series from the mean of vertices within two networks in standard-space Yeo17 atlas: “Yeo17-DNa” and “Yeo17-DNb.”

Within each run, EEG features were extracted across time, space, and frequency dimensions. We performed EEG time-frequency decomposition with complex Morlet wavelets applied to 40 bins (1-Hz spacing) for frequencies from 1-40 Hz. The number of wavelet cycles was scaled proportionally with the frequency (number of cycles = 0.5\**f*) to ensure an approximately constant temporal window across the frequency spectrum. The resulting EEG-BLP amplitude values (after wavelet filtering) were normalized over time within frequency bands and channels; to do so, we divided power at each time point by the total power across all time points within each frequency band and channel. We then downsampled the EEG-BLP timeseries to match the fMRI resolution (0.5 Hz) by applying an anti-aliasing fourth-order Butterworth low-pass filter, as has been done in prior work,^45,115^ yielding one zero-lag EEG-BLP estimate (per channel and frequency) aligned to the onset of each fMRI TR. To reduce dimensionality of EEG-BLP features and for the purpose of statistical inferences, we performed averaging across all channels. This channel-wise averaging step allowed simplified statistical testing and was further motivated by our attempt to replicate prior findings which indicated that EEG-BLP correlates of BOLD activity in the canonical DN were based on spatially diffuse EEG signals.^34,36,45^

We computed temporal correlations (Spearman’s ρ) between each EEG-BLP estimate (averaged across channels) and BOLD time series for Yeo7-DN, DNa and DNb. These correlations were computed at multiple temporal offsets between EEG-BLP and BOLD to account for the expected hemodynamic delay of the BOLD signal relative to neurophysiological activity.^15,116^ In particular, lags of 0 to -20 seconds were included (in increments of 2 seconds). Lags between -2 to -10 seconds are often considered to be physiologically relevant,^60^ but relationships between EEG-BLP and BOLD at shorter delays (0 to -2 seconds) have been reported.^40,45^ Our inclusion of longer delays (-10 to -20 seconds) allowed for specificity analyses. Excluded from all EEG-BLP correlation analyses were the first 10 BOLD samples in each run (i.e., to avoid inclusion of lag-shifted EEG samples acquired prior to fMRI onset) as well as samples that included high head motion (censored, as specified above). We elected not to convolve EEG-BLP measures with a hemodynamic response function (HRF), given that data-driven approach already considered multiple lags and that the HRF shape varies across individuals, brain regions, and electrophysiological frequencies of activity.^40,117^

We statistically compared the magnitude of EEG-BLP correlation with DNa versus DNb at the group level based on a breakdown of spatially-averaged EEG features into canonical frequency bands (similar to prior work^45^). We averaged EEG-BLP within the following bands: delta (1-3 Hz), theta (4-7 Hz), alpha (8-12 Hz), beta1 (13-20 Hz), beta2 (20-30 Hz), and gamma (31-40 Hz). The split of the beta range into two bands was motivated by prior work indicating the correlations between BOLD signals and EEG-BLP may be variable across higher versus lower beta ranges.^33^ Similar to Xavier et al.,^45^ we computed lag-averaged EEG-BLP values across the 0 to -10 second window (with 2-second bins) prior to each fMRI TR given the relevance of this lag range to the hemodynamic delay. Using these lag-averaged EEG-BLP values, we computed temporal correlations (Spearman’s ρ) with DNa and DNb BOLD activity within each run, performed Fisher *z* transformations on the ρ values, and then averaged the *z* values across all runs within each subject. We submitted the *z* values (representing correlation between EEG-BLP and BOLD activity) to a two-factor, within-subject repeated measures analysis of variance (two-sided), where factors were frequency band (delta, theta, alpha, beta1, beta2, gamma) and network (DNa, DNb). We implemented false discovery rate (FDR) corrected post-hoc Wilcoxon sign-rank tests (threshold of *p*=0.05) to compare the two networks in terms of EEG-BLP correlation with BOLD activity at each frequency band. We repeated the same analyses for personally-mapped DNa versus Yeo17-DNa and personally-mapped DNb versus Yeo17-DNb.

To further test the spatial specificity of DNa versus DNb differences in correlations with EEG-BLP, we conducted a spatially-constrained null model analysis.^118^ For this analysis, we constructed a null model in which we took the 43 original parcels that were labeled as DNa (23 parcels) or DNb (20 parcels) from the output of areal MSHBM,^69^ as described above. Across 1000 permutations, we randomly assigned the network identity of each of the 43 parcels while maintaining the total number of parcels per network. This resulted in 1000 versions of “surrogate a” (23 parcels) and “surrogate b” (20 parcels) networks that each included spatially interdigitated regions (i.e., similar to the original DNa and DNb) but with random organization within the common territory comprising DNa and DNb. Within each permutation, we extracted the mean BOLD time series from the surrogate networks. We then computed temporal correlations (Spearman’s ρ) with the EEG-BLP features used in the DNa versus DNb analysis described above (i.e., channel-averaged and lag-averaged EEG-BLP within six frequency bands). Next, we computed the differences of the correlations (surrogate network “a” minus surrogate network “b”) between the two surrogate networks. For each permutation, the inter-network difference was computed within individuals (averaged across runs then across session), and then a group-level average was computed across individuals. The 1000 group-level average values (i.e., the null distribution) obtained from all permutations was then compared to the observed group-level average difference between DNa and DNb for each of the six frequency bands. An FDR-corrected *p* value was computed for each frequency based on the rank of the observed effect relative to the null distribution.

While primary analyses were based on spatially-averaged EEG-BLP across electrodes, we further explored whether correlations were spatially heterogenous. We computed channel-wise temporal correlations between EEG-BLP (lag-averaged between 0 to -10 seconds of EEG-BLP relative to BOLD) and averaged BOLD activity within Yeo7-DN, DNa, and DNb. We plotted topographic scalp maps (averaged within canonical frequency bands) as well as channel-wise correlation heatmaps for Yeo7-DN, DNa, and DNb. Importantly, alpha (∼10 Hz) oscillations have been linked to the classical visual ‘idling’ rhythm at posterior regions but to sensorimotor rhythms (termed “mu” rhythm) at pericentral regions.^119^ To account for this factor, we generated DNa and DNb correlation heatmaps that were restricted to either posterior occipital channels (average of O1, O2, and OZ)^33,37,120^ or pericentral channels (average of C3 and C4).^119^

Beyond the aforementioned analyses that were specifically focused on DN subdivisions, we conducted wider mass-univariate analyses of all six networks of interest and compared pairs of parallel networks (i.e., DNa vs. DNb, FPCNa vs. FPCNb, and dATNa vs. dATNb). The EEG features that we included in these mass-univariate analyses were EEG-BLP, averaged within six canonical frequency bands (delta, theta, alpha, beta1, beta2, and gamma), at six delays relative BOLD sampling (0, 2, 4, 6, 8 and 10 seconds). We performed mass-univariate correlation (Spearman’s ρ) between the resulting 32 EEG-BLP features and BOLD activity within each network. The *ρ* values were Fisher *z*-transformed and averaged across all runs within each subject. For each network, statistical significance was assessed at the group level for each EEG-BLP feature via sign tests for zero median. The magnitude of correlation with each EEG-BLP feature was further compared between parallel network pairs using two-sided Wilcoxon signed rank tests. For all of these mass-univariate analyses, FDR correction was applied at the network level with a threshold of *p*=0.05 to correct for multiple comparisons.

### EEG-fMRI predictive modeling analyses

We conducted machine learning analyses to determine whether BOLD activity within each network of interest (DNa, DNb, FPCNa, FPCNb, dATNa, dATNb) could be modeled and predicted from EEG features. Given the ability of predictive models to handle large feature sets, EEG data were processed using a pipeline distinct from the mass-univariate analyses to ensure retention of a broader range of features, in line with prior work involving machine learning to predict fMRI from EEG.^40,115,121^ Time–frequency decomposition was performed using Morlet wavelets to estimate power amplitude at frequencies spanning 1–40 Hz, sampled at 1-Hz resolution. Prior to each BOLD TR marker, power amplitude estimates were extracted from 20-s intervals and organized into non-overlapping 2-s time bins, yielding 10 time bins per BOLD sample. Power values were then log-transformed and z-score normalized within subjects to zero mean and unit variance. This normalization was performed independently for each EEG channel and frequency across all samples and time bins for a given subject and was not performed separately by run or session. Frequencies were subsequently averaged into canonical bands, including delta (1–4 Hz), theta (4–8 Hz), alpha (8–13 Hz), beta (13–30 Hz), and gamma (30–40 Hz). Normalized power values were averaged within each frequency band and time bin, resulting in feature tensors of dimensions channels × frequency bands × time bins for each BOLD sample. For our main model, channel-wise features were retained, but we also conducted a comparison analysis where features were averaged along the channel dimension (i.e., resulting in spatially global features). As our predictive modeling approaches included minor divergences from the univariate analysis pipeline (e.g. in precise frequency band definitions) and considered joint information across features, this imposed some limits on making direct comparisons between univariate and machine learning results.

Prediction of BOLD activity from each network from EEG features was performed using XGBoost regression, a gradient-boosted decision tree method.^63^ Gradient-boosted decision trees are an ensemble-based optimization framework in which a sequence of weak learners—typically shallow decision trees—are combined to form a strong predictive model (**Fig. S12**). The process begins with an initial weak base learner that provides a coarse functional approximation. Gradient boosting then proceeds in a stage-wise additive manner, where each subsequent tree is trained to fit the negative gradients (i.e., the residual errors) of the current ensemble with respect to a chosen loss function. These residuals represent the discrepancy between predicted and true values and serve as pseudo-targets for the next tree. By iteratively fitting weak learners to these gradients and updating the model, the algorithm performs functional gradient descent, gradually refining the approximation. Through this accumulation of weak learners, the ensemble converges into a high-accuracy strong learner.

We selected the XGBoost method because of its ability to capture nonlinear relationships and multivariate feature interactions that may not have been detected in mass-univariate analyses. Moreover, XGBoost can effectively handle high-dimensional and correlated predictors, while model features can be interpreted using Shapley additive explanations (SHAP). Separate models were trained for each network using fixed hyperparameters (300 trees, maximum tree depth of 3, learning rate of 0.05, subsampling ratio of 0.8, column subsampling ratio of 0.8, L2 regularization λ = 1, squared-error loss). Cross-validation was performed using a leave-one-subject-out framework, in which data from one participant (including both recording sessions) were held out for testing while models were trained on data from all remaining participants. Importantly, EEG features were normalized at the subject level prior to cross-validation, ensuring that no information leakage occurred between training and test sets. Prediction accuracy was quantified using the coefficient of determination (R²), computed separately for each subject and network. Statistical inference on prediction performance was conducted using one-sample, one-sided Wilcoxon signed-rank tests against zero for each network, with multiple comparisons controlled using the Benjamini–Hochberg false discovery rate procedure.

To interpret the EEG features contributing to the prediction of BOLD activity within each network, we refit XGBoost models on all valid samples and computed SHAP values to decompose each model’s predictions into contributions from individual EEG features, defined by channel, frequency band, and time bin. Positive and negative SHAP values indicate whether a given EEG feature increases or decreases the predicted BOLD value relative to the model’s expected (baseline) prediction. Feature importance was summarized by computing mean absolute SHAP values. The SHAP values were first averaged across trials within each subject and then averaged across subjects to obtain group-level importance estimates. Statistical significance of SHAP-derived feature importance was assessed using within-subject permutation tests applied to reduced summary representations. Specifically, permutation testing was performed on band × time maps (after averaging |SHAP| across channels) and on channel × band maps (after averaging |SHAP| across time bins). For each permutation (n = 500), BOLD values were randomly permuted within subjects, preserving subject-level structure while disrupting the temporal correspondence between EEG features and BOLD signals. For each permuted dataset, XGBoost models were refit and SHAP values recomputed, yielding empirical null distributions of mean absolute SHAP values. Empirical p-values were computed by comparing observed SHAP values to the permutation-based null distributions. Multiple comparisons within each summary map (band × time and channel × band) were controlled using the Benjamini–Hochberg false discovery rate (FDR) procedure. To further enhance interpretability of results, we plotted spatial variability of |SHAP| values on scalp-level topographic maps where one map was generated per network and frequency band (averaged across time bins).

### Inter-session reliability analyses

We explored the reliability of EEG-BLP correlations with fMRI networks via comparing independent data from session #1 versus #2. We repeated the mass-univariate analyses described above except that we performed averaging at the session level within each subject. For all EEG channels and lags of EEG relative BOLD sampling (ranging from 0 to -20 seconds in increments of 2 seconds), we computed the intraclass correlation coefficient (ICC; two-way mixed, single measures, consistency) for session #1 versus #2. Each ICC value thus represented the degree to which temporal correlation between a given EEG-BLP feature and an fMRI network was consistent within subjects between sessions. Concatenating the ICC for all correlations with the six target fMRI networks, we classified ICC for each EEG-BLP feature as poor (<0.40), fair (0.40-0.59), good (0.60-0.77), or excellent (0.75-1.00). We sorted these classifications by lag of EEG-BLP feature relative to BOLD sampling because we predicted that features computed at neurophysiologically relevant lags (i.e., 0-10 seconds) would show greater reliability than those computed at later lags (i.e., 12-20 seconds).

We further explored how reliability changed as a function of the amount of data used within each session to compute ICC. As the duration of runs varied between subjects (especially for the self-paced rs-MDES condition), we first trimmed all runs to the minimum duration within any given subject so that durations were matched across all datasets. This trimming resulted in ∼8 mins of data per run. We plotted mean ICC value across all EEG-BLP features (limited to lags of 0 to -10 second relative to BOLD sampling) as a function of number of runs included in computation (ranging from 1 to 4) and broken down by network. We further computed mean ICC values for all EEG-BLP features using the full dataset within each subject (i.e., without trimming runs) and identified the maximum ICC values for each network based on a screening of all EEG-BLP features.

### Task (GradCPT) versus resting state analyses

We examined whether mass-univariate associations between EEG-BLP and network-specific BOLD activity varied as a function of condition (GradCPT performance versus resting state). Due to acquisition errors during GradCPT data collection, three subjects had only one usable GradCPT run and were thus excluded from these analyses (i.e., 20 subjects were included). For these analyses, we deleted experience sampling periods from rs-MDES runs to ensure that EEG relationships with fMRI were constrained to windows involving spontaneous neural activity (rather than cognitive and motor processes during responses to thought probes)., We deleted all BOLD samples during experience sampling periods as well as 8 seconds of samples following the offset of the last thought probe within each period to account for recovery of the BOLD signal. Thus, the resting state EEG-fMRI data that were included for the purpose of comparison to GradCPT were limited to visual fixation windows. After further accounting for head motion censoring, we retained an average of 830 TRs per subject (i.e., 27.7 minutes) across two sessions for the resting state condition.

Our task and resting state analyses proceeded in the same manner as the mass-univariate analyses described above, except correlations between EEG-BLP and each fMRI network were performed independently for data from each condition. For each condition and network, statistical significance was assessed at the group level for each EEG-BLP feature via sign tests for zero median. We further compared the magnitude of correlation with each EEG-BLP feature betweenGradCPT versus rest for all networks using two-sided Wilcoxon signed rank tests. For all condition-specific and cross-condition mass-univariate analyses, FDR correction was applied at the network level with a threshold of *p*=0.05 to correct for multiple comparisons.

## Data Availability

Raw data and preprocessed EEG data are openly available at the OpenNeuro repository (https://openneuro.org/datasets/ds007216) and are co-deposited in the NIMH Data Archive (collection number C4336). Individualized network parcellations and BOLD time series are available from https://github.com/DynamicBrainMind/precisionEEGfMRI.

## Code Availability

Original code for analysis and modeling are available from https://github.com/DynamicBrainMind/precisionEEGfMRI.

## Acknowledgments

This work was supported by the National Institute of Mental Health of the National Institutes of Health under award number R21MH127384 to AK. The content is solely the responsibility of the authors and does not necessarily represent the official views of the National Institutes of Health. This material is also based upon work supported by the National Science Foundation Graduate Research Fellowship Program under Grant No DGE2439563 to LST. Any opinions, findings, and conclusions or recommendations expressed in this material are those of the author and do not necessarily reflect the views of the National Science Foundation. We thank Rodrigo Braga for providing comments on a draft of this manuscript.

## Author Contributions

AK: Writing – original draft, Writing – review and editing, Formal analysis, Conceptualization, Methodology, Supervision, Project Administration, Software, Visualization, Validation, Data Curation, Funding Acquisition.

DB: Writing – review and editing, Methodology, Formal analysis, Visualization, Software, Data Curation, Validation.

LST: Writing – original draft, Writing – review and editing, Investigation, Methodology, Formal analysis, Project administration, Data Curation, Visualization, Writing – review and editing, Investigation.

AA: Writing – original draft, Write – review and editing, Methodology.

DP: Writing – original draft, Writing – review and editing, Formal analysis, Methodology, Software, Visualization, Validation.

