## Supplementary Info for "Electroencephalographic activity patterns distinguish interdigitated default and frontoparietal networks"

Kucyi et al.

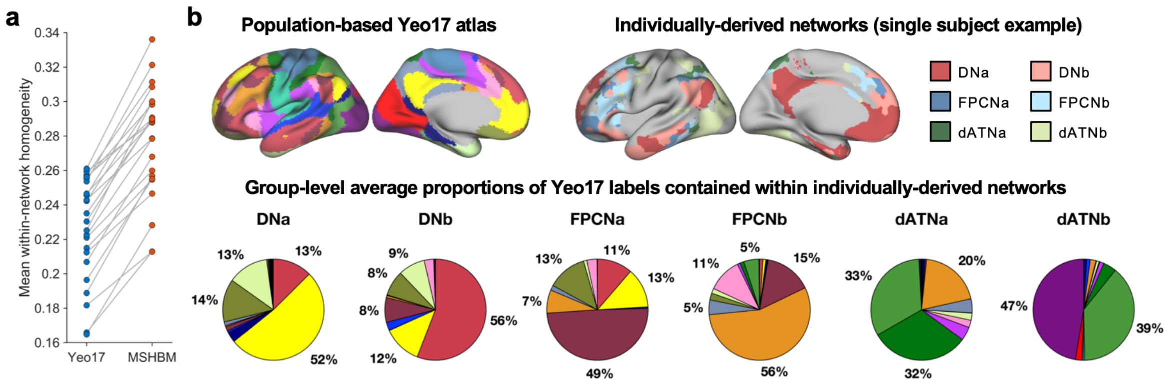


**Figure S1. Differences between networks derived from individual-level parcellation (using the MS-HBM method) versus a population-based atlas (Yeo17). a)** Homogeneity of functional connectivity profiles was computed for within-network vertices (see Methods). Homogeneity values were averaged across networks and sessions. As shown, for each subject (n=23), homogeneity was increased when networks were derived from MS-HBM relative to when derived from the Yeo17 atlas. **(b)** For each individually-derived network of interest (top right), labels from the population-based Yeo17 atlas (top left) were mapped onto the individualized parcellations. Pie charts show the cross-subject average proportions of vertices within each individually-derived network that overlap with each Yeo17-atlas network (each indicated as a unique color). The individually-derived DNa, DNb, FPCNa, and FPCNb show ~50% of their vertices mapping onto the expected corresponding Yeo17 networks. The individually-derived dATNa and dATNb show greater divergence from the Yeo17 atlas, showing more mixed contributions from different Yeo17 networks.

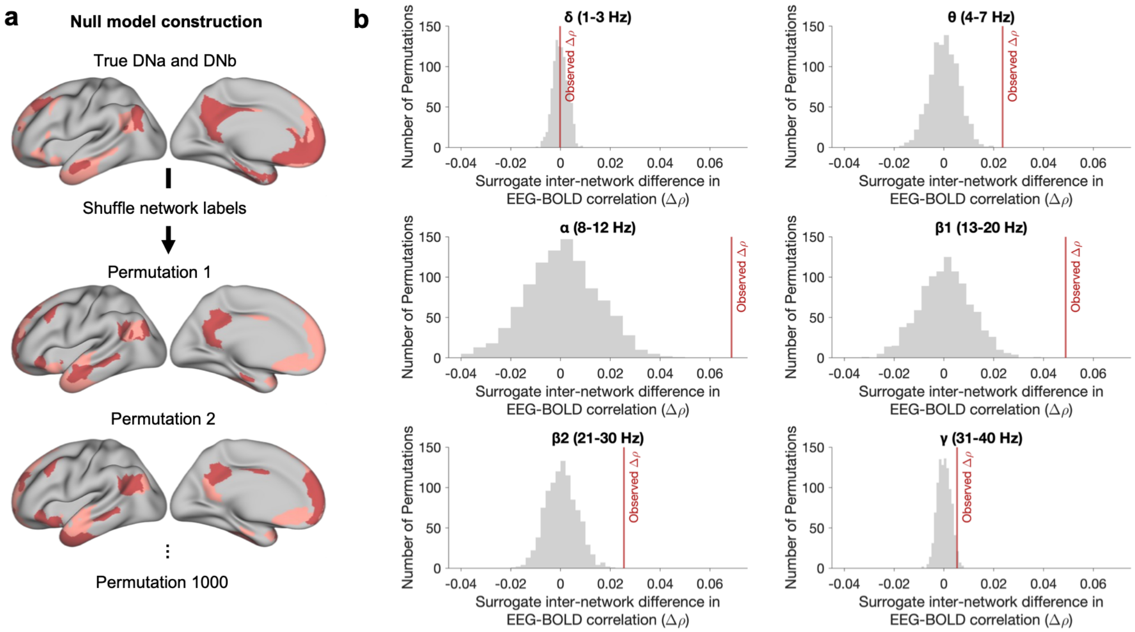


**Figure S2. Null model analysis for differences between DNa and DNb in coupling with EEG-BLP. a)** In each permutation (n=1000), parcels originally assigned to DNa and DNb (the "true" networks) were randomly shuffled to generate surrogate network "a" and surrogate network "b." After extracting the BOLD time series from each surrogate network and computing Spearman's correlations (ρ values) with EEG-BLP, the difference between networks (Δρ) was computed for each permutation to generate a null distribution. **b)** Null distributions (based on surrogate networks) and observed Δρ (based on the true DNa and DNb) are shown for each frequency band. The observed Δρ exceeded the null values at a statistically significant level for 5 out of 6 frequency bands (delta: $P_{FDR}$ = 1.0, theta: $P_{FDR}$ = 0.0015, alpha: $P_{FDR}$ = 0.0015, beta1: $P_{FDR}$ = 0.0015, beta2: $P_{FDR}$ = 0.0015, gamma: $P_{FDR}$ = 0.035).

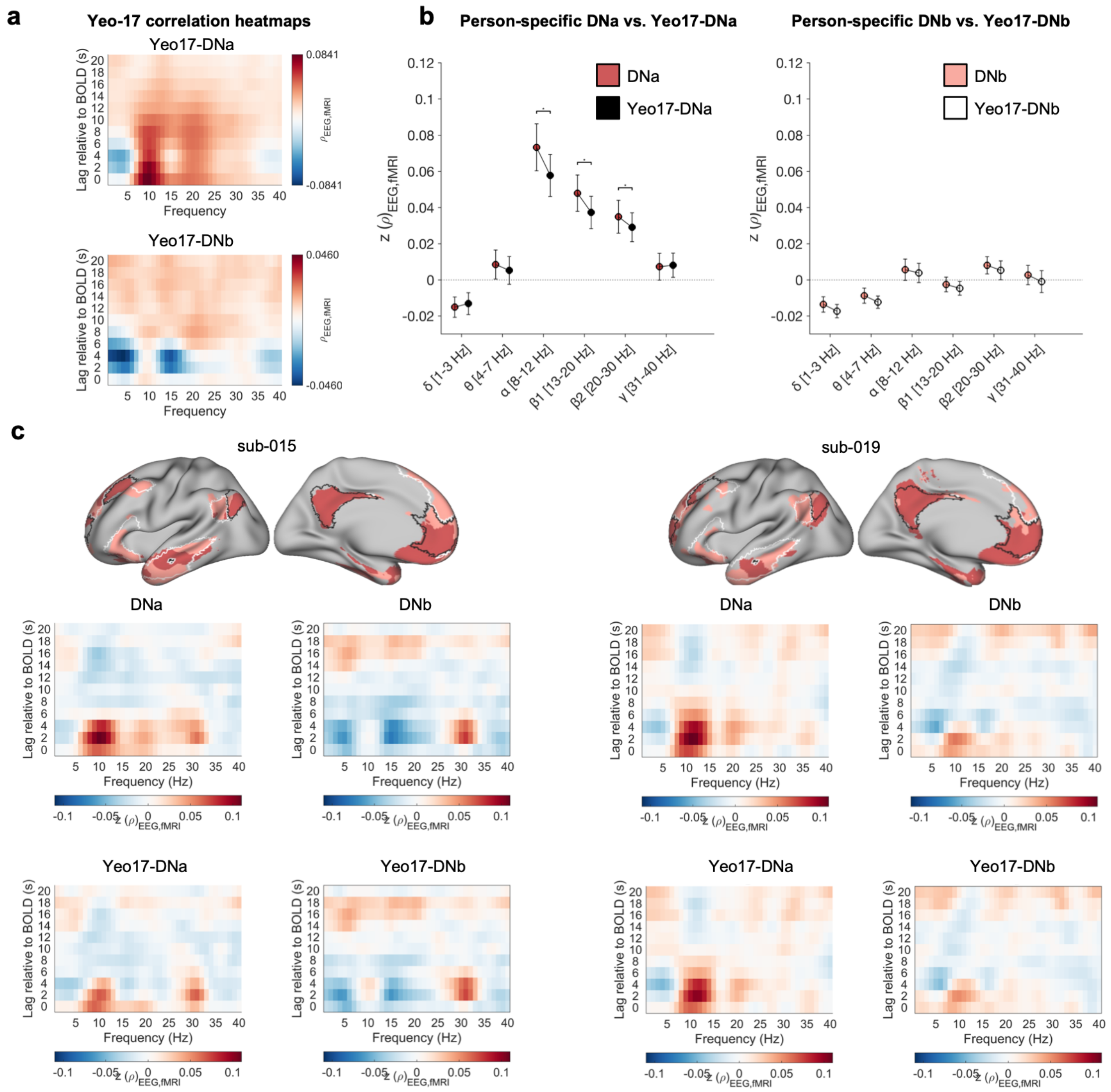


**Figure S3. Comparing personally-mapped DNa and DNb to population-derived versions of these networks in the Yeo 17-network atlas. a)** Spearman rank correlation (ρ) between EEG band-limited power (EEG-BLP) and BOLD activity at multiple lag shifts of EEG relative to BOLD data (average based on all subjects and sessions) for Yeo17-DNa and Yeo17-DNb. The EEG-BLP estimates were spatially averaged across all cortical electrodes. **b)** (Left) Comparisons between personally-mapped DNa (red dots) and Yeo17-DNa (black dots) and (Right) comparisons between personally-mapped DNb (pink dots) and Yeo17-DNb (white dots). The networks are compared in terms of temporal correlation with EEG-BLP at different frequency bands (spatially-averaged across electrodes and time-averaged for lags between 0 to -10 seconds relative to BOLD sampling). Correlations are shown as Fisher z-transformed ρ values based on all subjects and sessions. Black lines/asterisks indicate significant differences between networks in terms of the strength of BOLD association with EEG-BLP ($P_{FDR}$<0.05, Wilcoxon signed rank test). **c)** Examples of single-subject networks (top) and correlation heatmaps. On the cortical surfaces, black and white outlines, respectively, indicate Yeo17-DNa and Yeo17-DNb boundaries. Personally-mapped DNa and DNb, respectively, are shown in red and pink. An increased magnitude of EEG-BLP alpha correlation with DNa relative to Yeo17-DNa is observed in both subjects, which can be explained by misalignment between spatial locations.

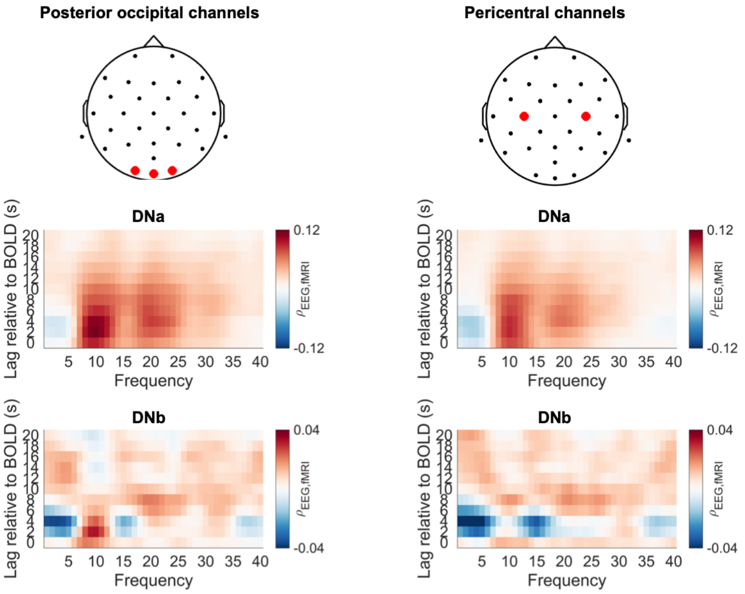


**Figure S4. Accounting for visual alpha versus sensorimotor mu in DNa and DNb correlations with EEG-BLP.** (Left) Correlation heatmaps based on EEG-BLP averaged from channels O1, O2, and Oz at posterior occipital locations. (Right) Correlation heatmaps based on EEG-BLP averaged from channels C3 and C4 at pericentral locations near sensorimotor cortex. The heatmaps display Spearman rank correlation (ρ) between EEG-BLP and BOLD activity at multiple lag shifts of EEG relative to BOLD data (average based on all subjects and sessions) for DNa and DNb. Alpha (~8-12 Hz) correlations with DNa appear as higher at posterior occipital relative to pericentral channels.

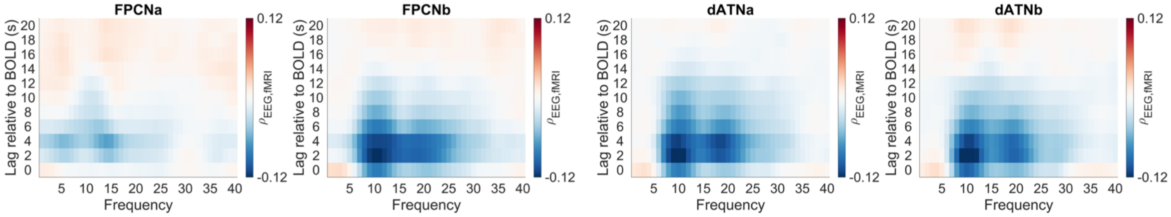


**Figure S5. Correlation heatmaps showing 1 Hz bins for EEG-BLP versus BOLD activity within FPCNs and dATNs.** Heatmaps show correlations between EEG-BLP and BOLD activity within each network. Correlations are shown as mean Spearman ρ values across subjects (n=23) for multiple frequencies and lags relative to BOLD (spatially-averaged across all EEG channels).

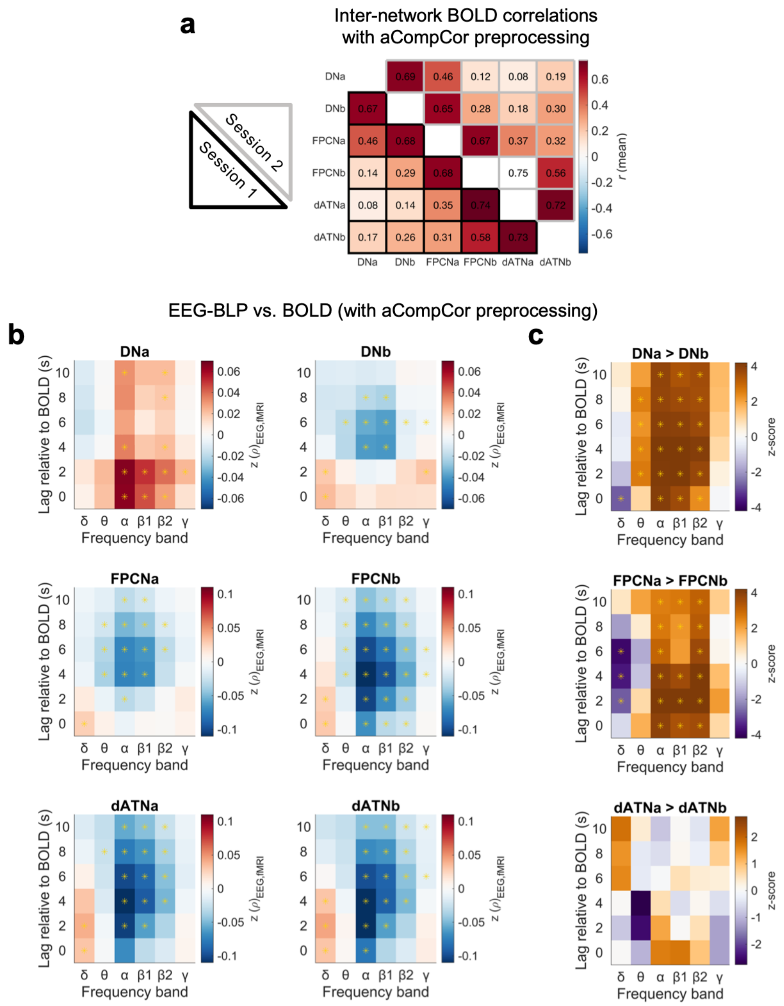


**Figure S6. Impact of aCompCor fMRI preprocessing on inter-network BOLD correlations and EEG-BLP versus BOLD correlations. a)** Group-level (n=23) mean temporal correlation values for BOLD signals between all pairs of networks (bottom left triangle: session #1, top right triangle: session #2). Values are based on averages across all fMRI runs within a given session when using the aCompCor preprocessing pipeline. See Fig. 1f for a comparison to results from the main global signal regression preprocessing pipeline. **b)** Correlation heatmaps showing EEG-BLP versus BOLD activity (based on aCompCor preprocessing) for each network, at lags ranging from 0 to -10 seconds, and for multiple frequency bands (based on all subjects and sessions). Associations are displayed as Fisher z-transformed Spearman rank correlation (ρ). Asterisks indicate significant correlations at the group level ($P_{FDR}$<0.05, sign test for zero median). See Fig. 3b for a comparison to results from the main global signal regression preprocessing pipeline. **c)** Comparisons (z-scores) between members of each parallel network pair for all frequencies and lags. Asterisks indicate significant differences at the group level ($P_{FDR}$<0.05, Wilcoxon signed rank test). See Fig. 3d for a comparison to results from the main global signal regression preprocessing pipeline.

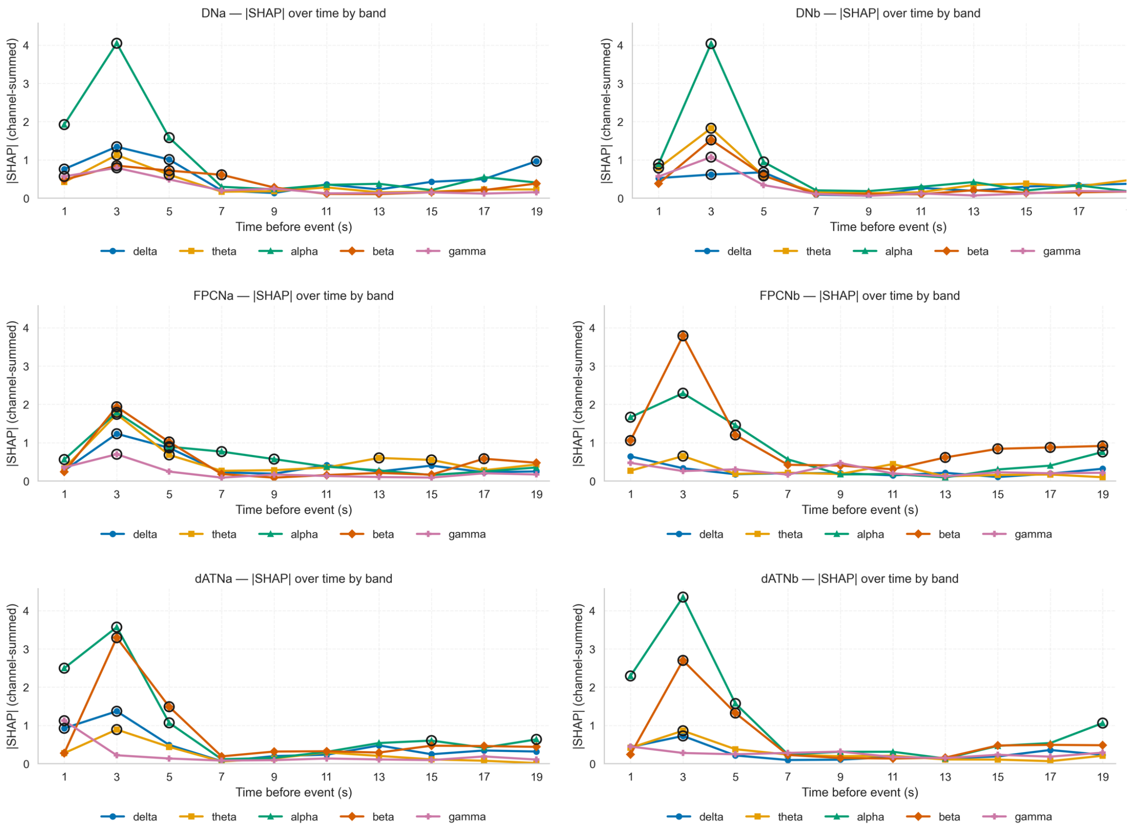


**Figure S7. Frequency-specific features that contributed to XGBoost predictive modeling of BOLD activity from EEG.** For each network, a plot displays the mean of absolute Shapley additive explanation (|SHAP|) values as a function EEG lag relative to BOLD activity ("Time before event"). Black circles indicate features that that showed significant contributions to predictive model performance ($p_{FDR}$<0.05, 500 permutations).

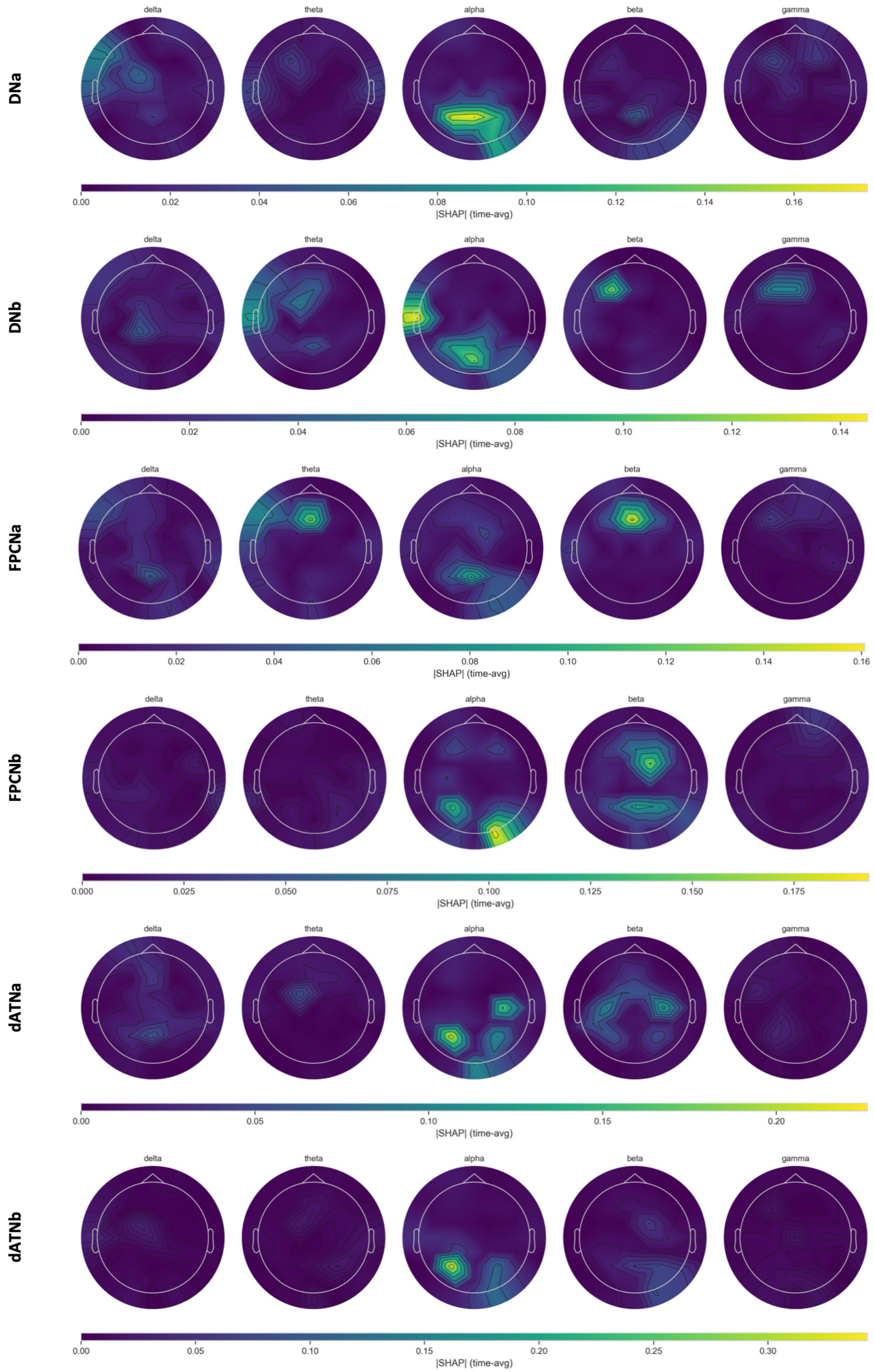


**Figure S8. Lag-averaged spatial features that contributed to XGBoost predictive modeling of BOLD activity from EEG.** For each network, topographic scalp maps display the mean of absolute Shapley additive explanation (|SHAP|) values as a function of frequency band. Yellow indicates spatial regions that contributed most strongly to XGBoost prediction performance. Prior to computing SHAP, spatial features were averaged across EEG lags relative to BOLD activity for the purposes of summary-level visualization of topographic distributions.

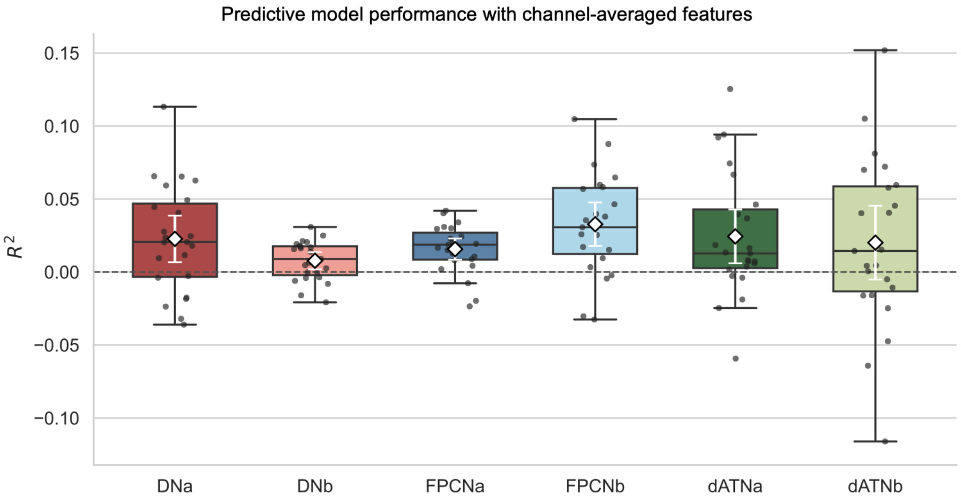


**Figure S9. XGBoost with spatially-averaged features (cf. Fig. 3e).** Results of predictive modeling using XGBoost and a leave-one-subject-out (LOSO) cross-validation scheme with channel-averaged features (i.e., the mean EEG-BLP for each frequency band across channels was used in modeling). Individual points show the coefficient of determination ($R^2$) for each held-out subject; the dashed horizontal line at $R^2$=0 indicates chance-level performance. Boxplots summarize the distribution of subject-level $\mathrm{R}^2$ values (center line: median; box: interquartile range, IQR = 25th–75th percentiles; whiskers: 1.5×IQR). White diamonds indicate the mean $R^2$ across subjects, with white error bars denoting the 95% confidence interval of the mean. For these channel-averaged predictive models, BOLD activity within all networks, except for dATNb, was successfully predicted from EEG-BLP features at an above-chance significance level (DNa: mean $R^2$ = 0.023, $p$ = 7.6x$10^{-3}$; DNb: mean $R^2$ = 0.008, $p$ = 8.9x$10^{-3}$; FPCNa: mean $R^2$ = 0.016, $p$ = 2.8x$10^{-4}$; FPCNb: mean $R^2$ = 0.033, $p$ = 1.1x$10^{-4}$; dATNa: mean $R^2$ = 0.024, $p$ = 3.4x$10^{-3}$; dATNb: mean $R^2$ = 0.020, $p$ = 7.1x$10^{-2}$). However, for each network, the model performance with channel-averaged features was weaker than it was when retaining features at the channel level (see Fig. 3e).

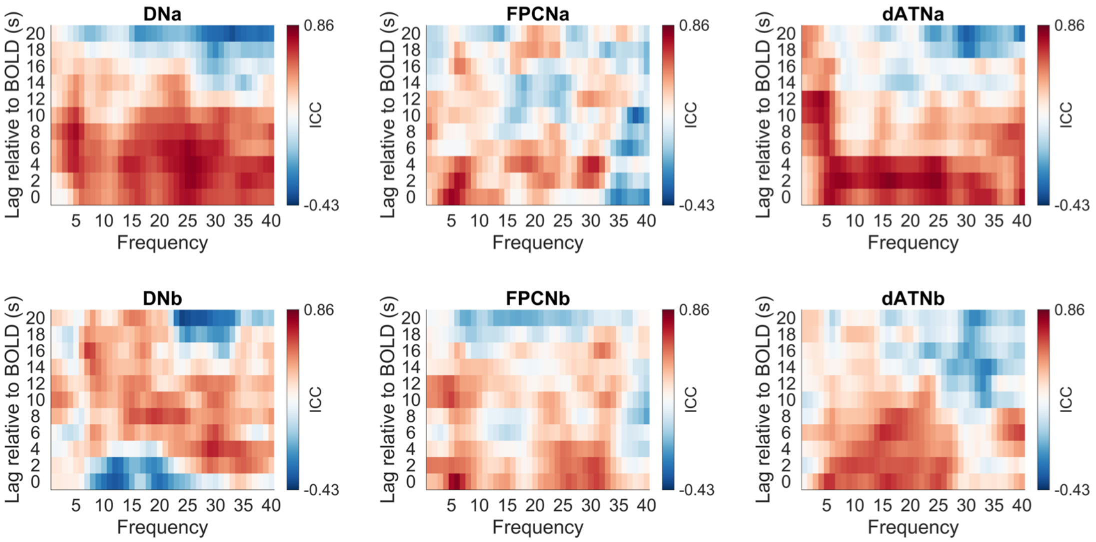


**Figure S10. Network-specific intra-class correlation maps for multiple frequency and lag pairings.** For each network, a plot displays intra-class correlation (ICC) values for EEG-BLP versus BOLD activity correlations estimated from session 1 versus session 2 data.

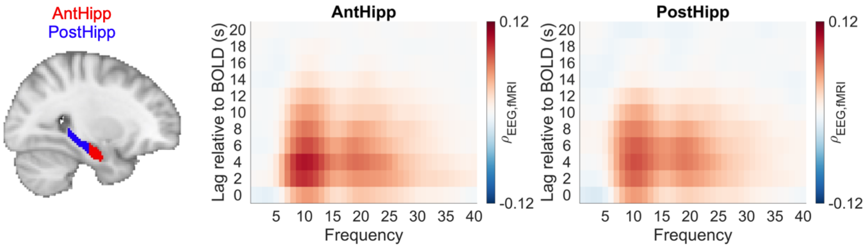


**Figure S11. Relationships between EEG-BLP and BOLD activity for anterior and posterior hippocampus.** The sagittal brain slice in MNI152 volumetric space shows the locations at which anterior and posterior hippocampal regions were defined. Heatmaps show correlations between EEG-BLP and BOLD activity within each hippocampal region. Correlations are shown as mean Spearman ρ values across subjects (n=23) for multiple frequencies and lags relative to BOLD (spatially-averaged across all EEG channels).

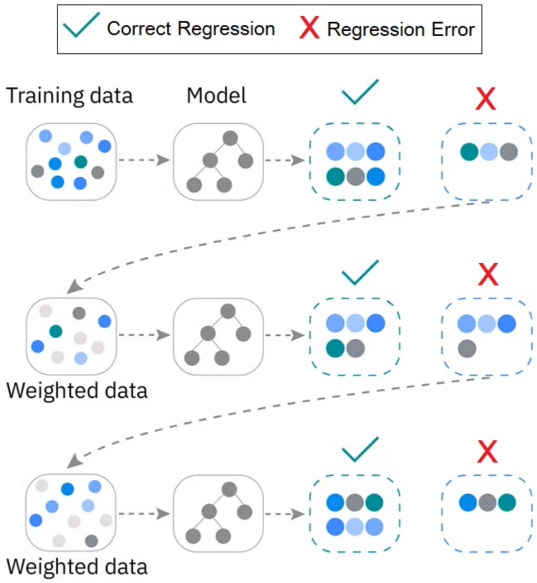


**Figure S12. Boosted tree ensemble model for the XGBoost method, combined from a sequence of weak decision trees.** Each weak tree is trained sequentially to correct the errors made by the previous trees. The final prediction for a given example is the sum of predictions from all trees, forming a strong regression model.
